# Structure of the human KEOPS/tRNA complex and characterization of pathogenic variants responsible for the Galloway Mowat syndrome

**DOI:** 10.64898/2026.03.03.709218

**Authors:** Charles Cirio, Sylvie Auxilien, Dominique Liger, Carlos A.H. Fernandes, Raoudha Dammak, Sophia Missoury, Emilie Zélie, Marion Dos Santos Malhao, Ana Andreea Arteni, David Touboul, Christelle Arrondel, Corinne Antignac, Géraldine Mollet, Catherine Vénien-Bryan, Herman van Tilbeurgh, Bruno Collinet

## Abstract

N^6^-threonyl-carbamoylation of adenosine 37 of ANN-type tRNAs (t^6^A) is a universal modification essential for translational accuracy and efficiency. The t^6^A pathway uses two sequentially acting enzymes, YRDC and OSGEP, the latter being a subunit of the multiprotein KEOPS complex. Structures of the subunits and subcomplexes of human KEOPS are known, but knowledge on the detailed interactions with tRNA is lacking. We present here the first structure of complete hKEOPS and of its complex with a substrate tRNA by cryo-electron microscopy. The CAA tail of tRNA is bound to the TPRKB subunit and the anti-codon loop is positioned at the entrance of the catalytic site of OSGEP subunit. The flexibility of the OSGEP-TP53RK interface allows hKEOPS to fit the surface of the tRNA elbow.

We recently identified mutations in all genes encoding for proteins of the t^6^A pathway in children with Galloway-Mowat syndrome (GAMOS), a clinically heterogeneous recessive disease characterized by early-onset steroid-resistant nephrotic syndrome and microcephaly. We here expressed and characterized the majority of the Galloway Mowat mutants. All mutants could be purified at high yields and seem to be stable *in vitro*. The t^6^A activity for most of the mutants is above 40% of the WT. Using CRISPR-Cas9 technology we replaced the genes encoding the t^6^A pathway proteins in yeast by their human homologues. This yeast construct was perfectly viable and produced WT levels of t^6^A modified tRNA. Using this tool, we observed that most of the GAMOS mutants were viable in yeast and yielded comparable t^6^A modified tRNA levels. Our data indicate that healthy human cellular development depends on an optimized level of t^6^A tRNA modification and is not compatible with a total loss of function of the t^6^A machinery.

## Introduction

Transfer RNAs (tRNAs) are composed of about 76 nucleotides and represent the adaptors between coding messenger RNA and nascent peptides on the ribosomes and thus perform a central function in the translation machinery. tRNAs are synthesized as precursors which, after trimming at the 5’ and 3’ends by dedicated nucleases, are chemically modified both at the bases and at the riboses at several positions (Gustilo et al., 2008; Väre et al., 2017). To date, about 150 tRNA modifications have been identified in bacteria, archaea and eukaryotes (Cappannini et al., 2024). Most modifications are located in the anticodon stem-loop region and have been shown to influence the accuracy and efficiency of the decoding process (Nedialkova and Leidel, 2015; Thiaville et al., 2016). Modifications outside this region often play a role in the stability of the tRNAs. The t^6^A modification is present in most ANN-decoding tRNAs and is the result of grafting a threonylcarbamoyl moiety onto the 6-amine of adenosine at position 37, immediately adjacent to the anticodon (Grosjean et al., 1995; El Yacoubi et al., 2011; Srinivasan et al., 2011; Perrochia et al., 2013a). The increase in translational fidelity in t^6^A-modified tRNAs was rationalized by structural studies (Beenstock et al., 2020). The presence of the threonylcarbamoyl group prevents the formation of a U_33_-A_37_ base pairing interaction and allows cross-strand stacking of A_38_ and t^6^A_37_ with the first position of the codon. The latter interaction enhances the codon anticodon base-pairing (Murphy et al., 2004). t^6^A is one of the rare universal tRNA modifications, which is reflected by the fact that its biosynthetic enzymes are present in virtually all sequenced genomes (Galperin, 2004; El Yacoubi et al., 2009). The biosynthesis of t^6^A modified tRNA occurs in two steps. The first is the synthesis of a chemically very unstable threonylcarbamoyl-adenylate (TC-AMP) intermediate from threonine, bicarbonate and ATP, which is catalyzed by the universal TsaC/YrdC/Sua5 family of enzymes (El Yacoubi et al., 2009). The threonylcarbamoyl moiety is subsequently transferred on the 6-aminogroup of A_37_ by the TsaD (bacteria)/ Kae1 (archaea, eukaryotes)/ Qri7 (mitochondria) enzymes (Wan et al., 2013; Suzuki and Suzuki, 2014; El Yacoubi et al., 2011; Srinivasan et al., 2011; Su et al., 2022). All these enzymes adopt a bi-lobal domain structure characteristic of the ASKHA (acetate and sugar kinases/Hsc70/actin) family of enzymes (Hecker et al., 2008; Mao et al., 2008; Nichols et al., 2013). In humans, the Kae1 homolog is named OSGEP and is a subunit of the multiprotein KEOPS complex (Arrondel et al., 2019). ATP or ADP bind in a cavity contained between the two lobes, carrying a metal that interacts with totally conserved histidines and the phosphates of the nucleotide (Hecker et al., 2008; Mao et al., 2008). Using a stable analogue of TC-AMP, it was shown for the bacterial TsaD that the intermediate occupies the same site as the nucleotides (Kopina et al., 2021). The other subunits of the human complex are GON7 (Gon7), LAGE3 (Pcc1), TP53RK (Bud32) and TPRKB (Cgi121), homologues in other organisms are bracketed (Downey et al., 2006; Kisseleva-Romanova et al., 2006; Wan et al., 2016; Arrondel et al., 2019). TP53RK is a small atypical protein kinase from the RIO kinase family that lacks the canonical activation loop. The role of its weak kinase activity, if any, has not been well established yet (Hecker et al., 2008; Mao et al., 2008). However in archaea, Bud32 has an ATPase activity that is important for the t^6^A activity of the KEOPS complex (Beenstock et al., 2022), but this activity has not been demonstrated for TP53RK. LAGE3 is a small three stranded β-sheet protein flanked by helices that interacts intensively with one of the OSGEP lobes. GON7 is an intrinsically disordered protein that becomes partially structured upon binding to LAGE3 (Arrondel et al., 2019). TPRKB is a single domain protein with a central anti-parallel β-sheet flanked by helices (Li et al., 2021). Recently cryo-EM structures of KEOPS complexes from *Arabidopsis* and archaea could be determined (Ona Chuquimarca et al., 2024; Zheng et al., 2024). The subunits in these complexes are arranged in a linear way with the order Pcc1, Kae1, Bud32 and Cgi121. A model of the human complex that was based on crystal structures of subcomplexes is in agreement with these structures (Arrondel et al., 2019). GON7, which is replaced by Pcc2 (a Pcc1 paralog) (Daugeron et al., 2023) in archaea and probably not yet identified in *Arabidopsis*, binds to LAGE3, opposite to the OSGEP subunit. The first detailed knowledge on the interaction with tRNA was raised by a recent study on an archaeal KEOPS complex showing that Cgi121, the TPRKB homologue, binds tRNA at the acceptor end and interacts with the CCA-tail (Ona Chuquimarca et al., 2024). The anticodon-loop is bound at the entrance of active site of the Kae1 enzyme in a non-active conformation. In this structure KEOPS adopts an elongated rigid binding platform whose shape is complementary to that of the tRNA elbow.

It is known since a few years that mutations in tRNA-modifying enzymes may lead to neurological and metabolic diseases (Suzuki, 2021; Zhang and Westhof, 2025). The Galloway-Mowat syndrome is a rare genetic disease characterized by an early onset nephrotic syndrome, microcephaly, brain anomalies and developmental delays (Galloway and Mowat, 1968). About ten different genes are now documented to be associated with GAMOS (Arrondel et al., 2019; Braun et al., 2017, 2018a; Fujita et al., 2018; Vodopiutz et al., 2015). Recessive mutations were identified in individuals with GAMOS within the *LAGE3*, *OSGEP*, *TP53RK*, and *TPRKB* genes (Braun et al., 2017). Later on, mutations were also detected in the *YRDC* gene in patients with a very severe form of GAMOS, suggesting that the t^6^A pathway is at the heart of the disease. Mutations in *GON7* lead to a milder form of the disease (Arrondel et al., 2019). Some of the cellular effects of the hKEOPS genes and their GAMOS mutations were investigated. Knockdown of OSGEP, TP53RK, or TPRKB inhibited cell proliferation, which could not be rescued by the GAMOS mutants (Braun et al., 2017). Considering that proteins involved in GAMOS concern the t^6^A pathway, it was not unexpected that impairment of these genes globally affected protein translation and caused endoplasmic reticulum stress. Despite these results, nothing is known about the effects of the GAMOS mutations on the t^6^A pathway at the molecular level.

In this paper, we characterized the structure of hKEOPS in complex with its tRNA substrate by cryo-electron microcopy at a resolution of 4 Å. In this complex TPRKB and TP53RK provide most of the interactions with the tRNA. The anticodon is bound near the active site groove of OSGEP in an inactive conformation. To better understand the effects of the molecular basis of the GAMOS, we set out to recombinantly express the majority of the documented GAMOS mutants and to characterize their *in vitro* biochemical properties, such as stability, t^6^A and ATPase activities and affinity for substrate tRNA. We further tested these mutants in a cellular context, by replacing all the yeast t^6^A pathway proteins by their human homologs using CRISPR-Cas9 technology and measuring the fitness and the capacity to synthetize t^6^A modified tRNAs of the mutant yeast strains. The purification of all the mutants led to stable hKEOPS complexes and the majority of these kept a t^6^A activity that was comparable to the WT proteins, both *in vitro* and in a yeast cellular context.

## RESULTS

### 1. Structure of hKEOPS bound to tRNA

To determine the structure of hKEOPS bound to substrate tRNA^Ile^_AAU_ we initially attempted crystallization of the complex without success and then turned to cryo-electron microscopy. The complex was first reconstituted by mixing recombinantly expressed hKEOPS subunits with *in vitro* transcribed tRNA^Ile^_AAU_ and then purified by gel filtration. Two experimental conditions proved to be essential for obtaining cryo-EM particles of the hKEOPS/tRNA^Ile^_AAU_ complex. First, we found that lowering the salt concentration in the purification buffer significantly increased the affinity of hKEOPS for fluorescently labeled tRNA (Fig. S1). A buffer containing 50 mM NaCl proved to be a good compromise between maintaining high hKEOPS-tRNA^Ile^_AAU_ affinity (Kd 54 +/− 3.4 nM) and avoiding aggregated particles during the preparation of the cryo-EM grids. Secondly, the best cryo-EM images were obtained with samples taken directly from the top-fraction of the gel filtration peak which contained the hKEOPS/tRNA^Ile^_AAU_ complex without additional concentration steps.

hKEOPS/tRNA^Ile^_AAU_ complex pure samples obtained from size exclusion chromatography were vitrified for cryo-EM imaging. Following 2D classification of the picked particles, the *ab-initio* reconstruction generated four different volumes of varying sizes. Besides, the expected volume corresponding to the hKEOPS complex bound to tRNA, three additional volumes were identified: the hKEOPS complex without the tRNA (apo-hKEOPS), isolated tRNA and, unexpectedly, a hKEOPS sub-complex bound to tRNA. Heterogeneous refinement was applied to the three volumes containing the hKEOPS complex, followed by non-uniform refinement steps of each independent volume, which yielded final high-resolution cryo-EM maps: 3.9 Å for the hKEOPS-tRNA complex, 3.7 Å for apo-hKEOPS, and 4.2 Å for the hKEOPS sub-complex bound to tRNA (Fig. 1 and S2). Significant local variation in resolution was observed across the cryo-EM maps. The highest resolution values were found in the core regions of the maps (3 Å) but resolution gradually declined towards the periphery of the complex, where resolution values ranged between 7 and 11 Å. Notably, the hKEOPS subunits exhibited higher resolution compared to tRNA (Fig. S2). The high-resolution cryo-EM data for the hKEOPS subunits enabled the construction of atomic models, guided by previously published crystal structures of the human hKEOPS subunits OSGEP-LAGE3-GON7 ((PDB ID 6GWJ (Arrondel et al., 2019)) and TPRKB-TP53RK ((PDB ID 6WQX (Li et al., 2021)), as well as the tRNA ((PDB ID 6UGG (Chan et al., 2020)). The analysis of the cryo-EM images revealed the presence of three types of particles: 39% corresponded to the hKEOPS/tRNA^Ile^_AAU_ complex, 39% to apo-hKEOPS and 22% to a hKEOPS/tRNA^Ile^_AAU_ subcomplex. However, due to the limited resolution in some regions corresponding to the tRNA and the peripheral areas of the complex, we could not unambiguously model the position of all tRNA bases and some protein side chains in the most peripheral areas of the complex.

**Figure 1.**
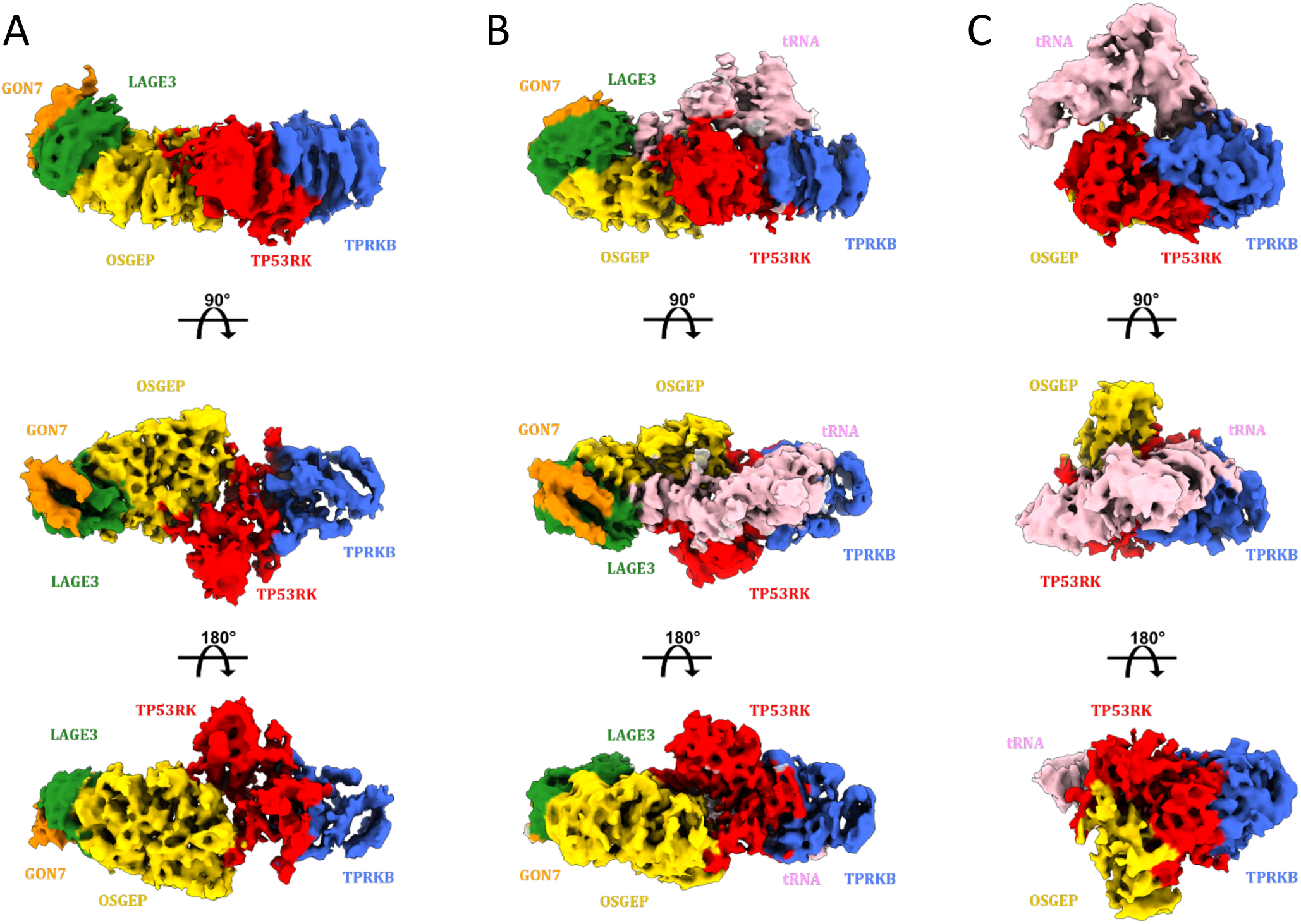
Cryo-EM maps of apo-hKEOPS, hKEOPS/tRNA, and hKEOPS/tRNA sub-complex. Subunits are colored as follows: TPRKB (blue), TP53RK (red), OSGEP (yellow), LAGE3 (green), GON7 (orange) and tRNA^Ile^_AAU_ is shown in pink. (A) Sharpened cryo-EM map of pentameric apo-hKEOPS, resolved at 3.74 Å. (B) Sharpened cryo-EM map of the hKEOPS/tRNA^Ile^_AAU_, resolved at 3.9 Å. (C) Sharpened cryo-EM map of a hKEOPS/tRNA^Ile^_AAU_ sub-complex resolved at 4.22 Å.

For both apo-hKEOPS and hKEOPS/tRNA^Ile^_AAU_, the five protein subunits were clearly identified, providing the first structures of a complete hKEOPS complex. Since high-resolution crystal structures were reported for all subunits, we focus our analysis on the architecture and flexibility of hKEOPS and its interaction with the tRNA^Ile^_AAU_ substrate.

As for archaeal and plant KEOPS, hKEOPS has a linear arrangement of its subunits (Ona Chuquimarca et al., 2024; Zheng et al., 2024). tRNA^Ile^_AAU_ bound to hKEOPS adopts a canonical L-shape and presents the inside of its elbow to the protein surface. Apart from GON7, all subunits of hKEOPS establish contacts with the tRNA. The quality of the maps was heterogenous and therefore difficult to interpret in some regions such as the anticodon loop and the T-arm of the tRNA^Ile^_AAU_. The T-loop (poorly defined in the maps) does not interact with hKEOPS (Fig. 2). tRNA^Ile^_AAU_ has its CCA acceptor tail positioned in a groove on the TPRKB surface and its AC-stem-loop at the entrance of the active site of OSGEP (Fig. 2). This binding mode of tRNA to hKEOPS is very similar to that observed in the recent cryo-EM structure of an archaeal KEOPS-tRNA complex (Ona Chuquimarca et al., 2024). However, in the human structure the CCA tail adopts a different orientation compared to the CCA tail in the archaeal complex (Fig. S3A). A_76_^tRNA^ is sandwiched between T74^TPRKB^ and Y65^TPRKB^ and makes a H-bond with the E112^TPRKB^ carboxylate while C_75_^tRNA^ stacks against R73^TPRKB^ and C_74_^tRNA^. TPRKB interacts also with the phosphate backbone of the tRNA acceptor stem region. The less well-defined cryo-EM map of the anticodon precludes a detailed description of its interactions with hKEOPS in this region. The anti-codon loop folds back slightly onto the stem region and is positioned between the C-terminal domains of TP53RK and OSGEP (Fig. S3B). It occupies the entrance of the active site groove of OSGEP without penetrating it. The backbone between nucleotides 30 and 35 makes a few contacts with LAGE3 and the backbone between nucleotides 27 to 29 interacts with the positively charged and well conserved C-terminal helix of TP53RK (Fig. S3C). In tRNA modifying enzymes, it is often observed that the target base of the tRNA pivots out of the core during the formation of the reactive complex. Although our map lacks details for the AC-region, we do not observe such a movement in our structure. Base A_37_ remains hidden in the core of the tRNA and makes a trans-anticodon contact with U33^tRNA^. Therefore, the 6-amino group of A_37_ is not available for modification in the present hKEOPS/tRNA^Ile^_AAU_ complex. In bacteria, the transfer of carbamoyl moiety is transferred by TsaB/TsaD/TsaE (TsaD is the bacterial homologue of OSGEP) (Zhang et al., 2015; Luthra et al., 2018; Missoury et al., 2018). To get better insight into the organization of the active site of OSGEP, we superimposed hKEOPS/tRNA^Ile^_AAU_ onto the crystal structure of the binary *E. coli* TsaB/TsaD complex bound to a stable TC-AMP analog (Kopina et al., 2021). This resulted in a very good structural superimposition between OSGEP and TsaD (RMSD of 1.5 Å) and the TC-AMP analogue could be accommodated in the active site of hKEOPS without any steric hindrance (Fig. S4). The overlay shows that the 6-amino group of A_37_ base is at a distance of 17 Å from the carbamoyl group of the TC-AMP analog. There is ample space in the active site of OSGEP to allow for a conformational change of the anticodon loop, which, combined with a pivoting movement of base A_37_ out of the tRNA core, should bring the 6-amino group into a favorable position for threonylcarbamoyl transfer. In the archaeal tRNA-KEOPS complex, a highly flexible anti-codon loop between position 33 and 39 was also observed (Ona Chuquimarca et al., 2024).

**Figure 2.**
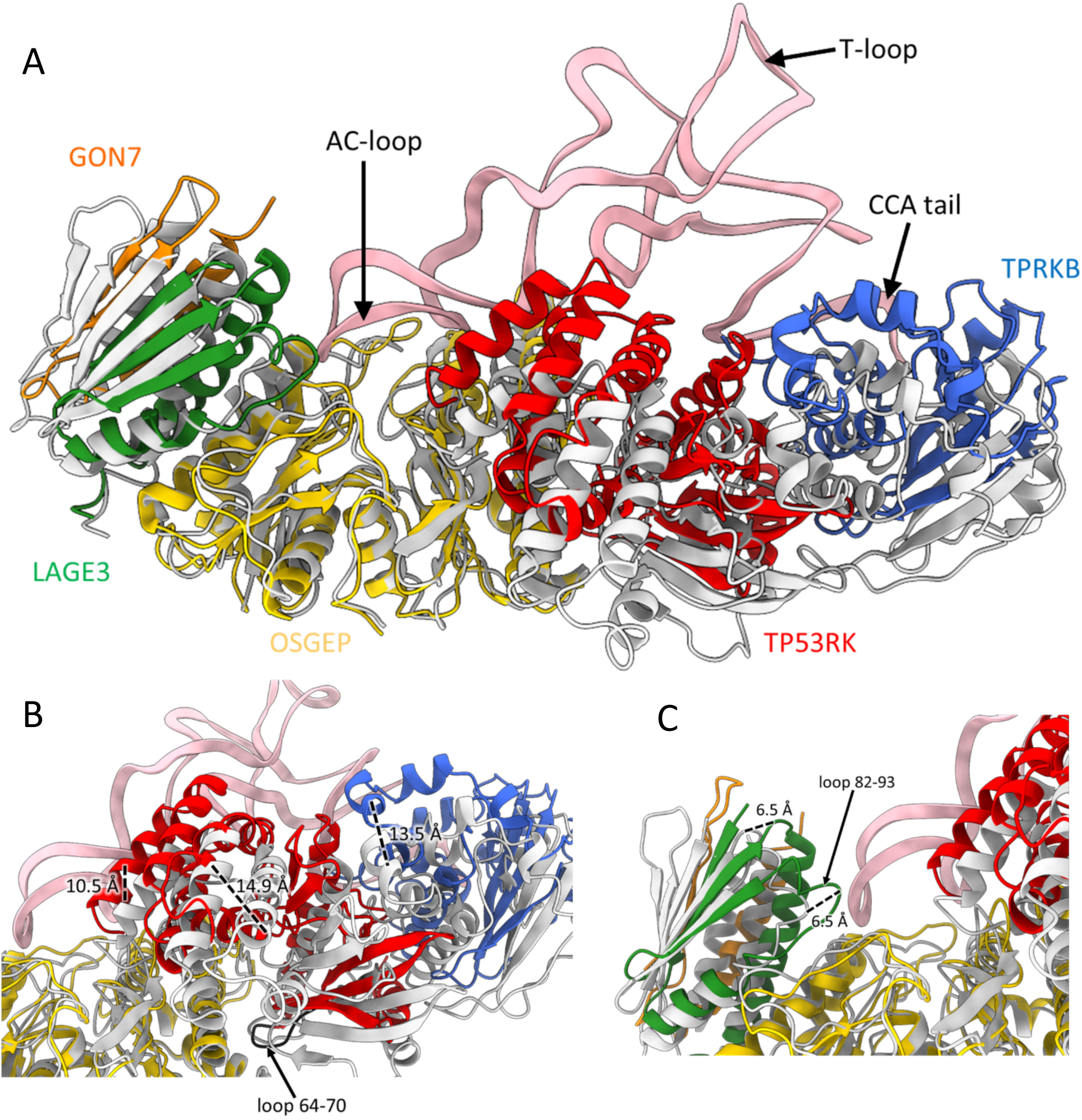
Superposition of apo-hKEOPS (grey) and hKEOPS/tRNA structures. Colored subunits are orange for GON7, green for LAGE3, yellow for OSGEP, red for TP53RK, blue for TPRKB and pink for tRNA^Ile^_AAU_. (A) Overall superposition reveals significant shifts of the TP53RK–TPRKB and LAGE3–GON7 subcomplexes upon tRNA binding. (B) Close-up of the TP53RK–TPRKB subunits highlighting the pivotal loop (residues 64–70), which shifts by more than 10 Å between the apo and tRNA-bound states but maintains a consistent contact between OSGEP and TP53RK (C) Close-up of the LAGE3–GON7 subunits showing loop 82–93 of LAGE3 positioned near the anticodon (AC) loop of the tRNA and an overall shifting of more than 6 Å

The phosphate backbone of G_10_C_11_ in the D-loop of tRNA is contacting conserved positively charged residues of TP53RK (K196^TP53RK^, K230^TP53RK^, K231^TP53RK^, R233^TP53RK^) which should electrostatically stabilize this interface. The critical role of the G_10_–C_25_ and C_11_–G_24_ base pairs for t^6^A biogenesis in hKEOPS was reported (Wang et al., 2022). Sequence alignments of tRNAs that activate the ATPase activity of archaeal Bud32 revealed the presence of a conserved C_11_U_12_ doublet that is absent in non-cognate tRNAs. Mutations of the bases in this region yielded tRNAs that were stable but could no longer be modified by hKEOPS (Wang et al., 2022).

The five subunits could be identified in the cryo-EM maps of apo-hKEOPS and the crystal structures of the GON7/LAGE3/OSGEP (PDB 6GWJ) and TP53RK/TPRKB (PDB 6WQX) subcomplexes fitted well into the maps (Fig. S5). Our apo-hKEOPS structure superposes very well onto the recently determined apo-AtKEOPS structure (RMSD of 1.6 Å) (Fig. S6A) (Zheng et al., 2024). To evaluate whether tRNA binding induces conformational changes into the hKEOPS complex we superimposed the cryo-EM structures of apo-hKEOPS and hKEOPS/tRNA^Ile^_AAU_. As shown in figure 2A, the OSGEP subunit matches well, but a considerable and complex movement of the TP53RK/TPRKB subunits is induced upon tRNA binding. The TP53RK/TPRKB subcomplex rotates around the loop contained between (P64-P70)^TP53RK^, which is one of the main contact sites between OSGEP and TP53RK in both apo and tRNA bound hKEOPS. This repositioning of TP53RK relative to OSGEP, creates a binding groove for tRNA and brings the TP53RK and TPRKB subunits into contact with the tRNA. For instance, the C-terminal helix of TP53RK shifts over about 11Å to establish contacts with the AC-stem (backbone of nucleotides 27 to 29) and the D-arm (nucleotides 9 and 10). These shifts also bring TPRKB into contact with the acceptor arm of the tRNA. The structures of the TP53RK/TPRKB subcomplex in apo- and hKEOPS/tRNA are very similar, suggesting that their movement towards the tRNA is principally caused by changes at the OSGEP-TP53RK interface (Fig. 2B). A similar 6 Å shift of the GON7/LAGE3 subcomplex is observed comparing apo- and tRNA-bound hKEOPS (Fig. 2C). The movement brings LAGE3 into contact with the tRNA backbone between nucleotides 31 and 35. We therefore conclude that the plasticity of the inter-subunit interfaces allows the hKEOPS complex to snuggle against the tRNA elbow. This flexibility was not observed in a recent cryo-EM study of archaeal KEOPS. In this case, tRNA binding did not seem to cause significant positional rearrangements of the KEOPS subunits. Comparison between the human and archaeal KEOPS (Fig. S6B-C), shows that the human OSGEP and archaeal Kae1 subunits superimpose well both in the apo- and the tRNA-bound forms. However, the position of archaeal Bud32/Cgi121 subcomplex relative to Kae1, is intermediate between that of the TP53RK/TPRKB subcomplex in the apo- and tRNA bound hKEOPS complexes.

For the archaeal complex, the presence of two tRNA conformers was reported, one exhibiting a native-like fold and the other had a distorted anti-codon domain conformation. The tRNA bound to hKEOPS, is rather similar to the native-like fold of the archaeal KEOPS bound tRNA. However, significant differences of the tRNA conformations between human and native-like archaeal complexes are observed at the level of the AC-stem loop (between nucleotides 28 and 38). It is known that the AC-loop regions of tRNAs have an inherent flexibility compared to the remainder of the elbow structure. Apparently, this flexibility is maintained even when bound to KEOPS. In the distorted conformation of the archaeal complex, the tRNA exhibited a flipping of the G_26_ base. This base is part of a network of interactions well conserved across KEOPS substrates. Mutations G_26_C and G_26_U in the archaeal tRNA enhanced t^6^A modification activity by a factor of 2 to 3, without significantly affecting tRNA affinity or ATPase activity. There was no evidence for G_26_ base flipping in hKEOPS/tRNA^Ile^_AAU._ However, TP53RK on the human complex interacts with the backbone of bases G_26_U_27_G_28_ and the phosphate establishing charged contacts with well conserved positively charged residues from the N-terminus (Fig. S7). It was further shown that the D-arm nucleotides are important for the activity of KEOPS (Wang et al., 2022).

About 20% of the cryo-EM particles correspond to an object that was considerably smaller than the discussed hKEOPS complexes. We were able to model TPRKB/TP53RK/tRNA^Ile^_AAU_ and residues 126–287 of the C-terminal domain of OSGEP into the density maps. However, we could not resolve the N-terminal domain of OSGEP or the C-terminal fragment encompassing residues 288–335. Additionally, the remaining LAGE3 and GON7 subunits of hKEOPS were not identified in the maps. We never noticed proteolytic susceptibility of OSGEP during protein purification, and it is highly unlikely that it became proteolyzed during the cryo-EM experiments. We therefore assume that the lack of density in the maps for one third of the complex must be ascribed to mobility. Although in this subcomplex, the tRNA^Ile^_AAU_ interacts with TPRKB in a similar manner as for the complete KEOPS, the association of tRNA^Ile^_AAU_ with TP53RK is different. The backbone of C_11_-U_12_-C_25_ and the anticodon stem of tRNA bind to a positively charged patch in the C-terminal helix of TP53RK (K230^TP53RK^, K231^TP53RK^, R233^TP53RK^, K237^TP53RK^). Superimposition of the hKEOPS/tRNA^Ile^_AAU_ or archeal KEOPS/tRNA^lys^_UUU_ complex onto the subcomplex shows that their tRNA conformations are very similar. However, the acceptor region (around nucleotide 70) is oriented slightly differently, causing dissociation between the AC-stem loop and OSGEP in the subcomplex (Fig. S8). We hypothesize this subcomplex could represent a precatalytic hKEOPS/tRNA complexes and that LAGE3-GON7 subunits could contribute to the positioning of the tRNA AC-loop in the active site of OSGEP.

The successful structure prediction program AlphaFold has recently acquired the capacity to model accurately structures of protein and nucleic acid complexes. We exploited the AlphaFold 3 version (AF3) to model the hKEOPS/tRNA^Ile^_AAU_ complex. As can be seen in figure S9, in the AF3 model of this complex, hKEOPS is very similar to our apo-cryo-EM structure (Fig. S9D, E). The acceptor arm of tRNA^Ile^_AAU_ is bound to TPRKB and the anticodon loop is inserted into the active site groove of OSGEP. tRNA adopts a canonical structure and its binding mode to KEOPS is very similar to that observed for the human and archaeal KEOPS tRNA-complexes. There is however a noteworthy difference of the conformations of the anticodon loop between the experimental and modelled structures. In the AF3 model the AC has an open conformation and most remarkably the A_37_ base has flipped out of the tRNA core (Fig. S9B). Superposition of the anticodon loop onto the model of a TC-AMP analog bound into the active site of OSGEP shows that the A_37_ base is now very close to the carbamoyl of the TC-AMP intermediate. A rotation around the C1’-N9 bond of A_37_ would bring the N^6^ amine group into position for a nucleophilic attack of the carbamoyl moiety (Fig. S9C).

### 2. Sampling of the hKEOPS/tRNA^Ile^_AAU_ interface

We set out to further explore the structure of the hKEOPS/tRNA^Ile^_AAU_ complex by mutating highly conserved residues at or near the hKEOPS-tRNA^Ile^_AAU_ interface, and measuring their t^6^A and ATPase activities, as well as their tRNA binding capacities. We restricted our selection to residues that had not already been mutated in previous published studies (Beenstock et al., 2020; Ona Chuquimarca et al., 2024). The positions of mutations are shown in figure S10 and biochemical data are gathered in figure S11. Single mutations that affect the t^6^A activity most severely are located in OSGEP. To be able to rationalize at best these mutations we used our model of the TC-AMP analogue bound to OSGEP (Fig. S4). In this model the phospho-threonyl part of TC-AMP is held between 2 short loops (residues 10 to 15 and 132 to 135). The Oγ of S10^OSGEP^ forms a hydrogen bond with the threonyl moiety and the carbamoyl group contacts A11^OSGEP^ (Fig. S12A). S10A^OSGEP^ has lost 60% of the t^6^A activity, while A11L^OSGEP^ and A11G^OSGEP^ are almost inactive. The phosphate of TC-AMP is bound both by K13^OSGEP^ and S132^OSGEP^ (Fig. S12A). K13N^OSGEP^ is inactive and S132A^OSGEP^ has lost 50% of its activity. The E181^OSGEP^ carboxylate is at 5.3 Å of the N^6^ amine of TC-AMP in the cryo-EM structure (Fig. S12B) but at hydrogen bond distance in the AF3 model. The E181A^OSGEP^ mutant has lost 70% of its t^6^A activity but retains its tRNA affinity. The Y130^OSGEP^ OH group provides one of the metal ligands in the active site, and Y130F^OSGEP^ displays a 30% decrease in t^6^A activity. This residue is conserved in eukaryotes and archaea but not in bacteria. Mutation of the two metal-bound histidines has previously been reported to completely abolish t^6^A activity (Kopina et al., 2021). None of the mutations around the TC-AMP binding site affect tRNA binding. Taken together, our mutational data strengthen the model of TC-AMP bound into the active site of OSGEP.

N159^OSGEP^ and R166^OSGEP^ are conserved residues of a Kae1 specific insert at the entrance of the active site groove (Ona Chuquimarca et al., 2024). Both contact the backbone of the AC-loop in our structure and in the AF3 model (Fig. S12C). Their mutation into alanine reduced activity by 60% and 80% respectively. The tRNA affinity of N159A^OSGEP^ was unaffected but decreased by 30% for R166A^OSGEP^. S205^OSGEP^ and S209^OSGEP^ are interacting with the D-arm in the AF3 model, but do not in our cryo-EM structure. Their mutation into alanine does not affect activity, and only slightly impacts tRNA binding affinity (Fig. S12E).

Next, we mutated a set of conserved positively charged residues of TP53RK. K238^TP53RK^, R245^TP53RK^ and R247^TP53RK^ are part of the positively charged C-terminal helix that interacts with the acceptor stem backbone. It was shown that the deletion of this helix causes loss of the t^6^A activity in archaeal KEOPS (Beenstock et al., 2020). R245^TP53RK^ interacts with the backbone of G_28_G_29_ and R247^TP53RK^ with the backbone of A_37_ (Fig. S12F). The t^6^A activities of R245Q^TP53RK^ and R247Q^TP53RK^ decrease by about 60% and the R245Q/R247Q^TP53RK^ double mutant is inactive. All these mutants also have weakened tRNA affinity. K238^TP53RK^ is at 4 Å from the U_27_ backbone phosphate. Its mutation into alanine weakens the tRNA binding but has only minor effect on t^6^A and ATPase activities. Together these data highlight the importance of the positively charged C-terminus of TP53RK for activity. R86^TP53RK^ is at 7 Å from the G_69_ phosphate in the cryo-EM structure and forms H-bonds with the phosphate backbone from nucleotides 69 and 70 in the AF3 model (Fig. S12F). The R86A^TP53RK^ mutant has lost 80% of its t^6^A and ATPase activity and diminished tRNA affinity. E194^TP53RK^ and D195^TP53RK^ are two highly conserved negatively charged residues that interact with the AC-arm backbone. Their mutation into alanine significantly increased the affinity for tRNA and caused only a minor effect on their t^6^A activity. Relieving the negative surface charge in this region therefore increases tRNA binding (Fig. S12G).

Mutation of three charged residues in TPRKB near the acceptor stem and CCA tail (R30A^TPRKB^, K147A^TPRKB^ and K150A^TPRKB^) (Fig. S12H) resulted in a 60% decrease in tRNA affinity (80% in the K147A/K150A^TPRKB^ double mutant) but had little effect on the t^6^A activity. Mutation of uncharged residues N88A^TPRKB^ and S90A^TPRKB^ had no effect. None of the TPRKB mutations affected t^6^A activity. These data suggest that electrostatic complementarity is a main contributor to tRNA binding affinity. Although the CCA tail of tRNA in our model has a different twist than that in the structure of MjKEOPS/tRNA, it binds to an equivalent region of TPRKB. The CCA tail of tRNA was shown to be essential for t^6^A activity of archaeal KEOPS. On the other hand, the *Pyrococcus abyssi* t^6^A synthesis can do without the Cgi121 protein (Perrochia et al., 2013b). Removal of the CCA end was shown to reduce the t^6^A efficiency of *S. cerevisiae* KEOPS and delete the t^6^A activity of both human and *C. elegans* KEOPS (Wang et al., 2022). Three mutants R73E^TPRKB^, I89E^TPRKB^ and L93E^TPRKB^ that were reported to have lost their affinity for tRNA (Beenstock et al., 2020), are in contact with the CCA tail in our structure (Fig. S12D). In summary, despite a slightly different orientation of the CCA tail, our cryo-EM data are in good agreement with the Cgi121-tRNA structure and confirm the mutational data from TPRKB.

### 3. In vitro characterization of GAMOS pathogenic variants

GAMOS mutations were identified in all five subunits of hKEOPS (reviewed in (Zhang and Westhof, 2025)), making it challenging to formulate an *a priori* hypothesis about the molecular mechanisms underlying this severe disease. To gain insight, we conducted an extensive biochemical characterization of near all documented GAMOS missense pathogenic variants. First, we overexpressed 32 mutant hKEOPS complexes in *E. coli*, following the same procedure used for the wild-type (WT) construct (see supplementary methods). All mutants were successfully purified to homogeneity and concentrated to at least 1 mg/mL. Their gel filtration profiles were identical to those of WT hKEOPS, indicating that the GAMOS mutations do not disrupt the folding or structural integrity of the recombinant hKEOPS complex (Fig. S13). In figure 3, we mapped the GAMOS mutations onto the 3D structure of the hKEOPS/tRNA complex, revealing that they are spatially scattered across four subunits of hKEOPS. Notably, the GAMOS mutation in GON7 results in a complete loss of the protein. To further interpret the biochemical properties of the GAMOS mutants, we used our model of the hKEOPS/tRNA/TC-AMP complex (Fig. S4). Interestingly, none of the GAMOS mutations directly impact inter-subunit interactions. This is consistent with our gel filtration data, which indicated that the quaternary structure is preserved in all expressed mutants. To identify potential molecular defects caused by the GAMOS mutations, we measured their t^6^A and ATPase activities as well as their affinities for tRNA. As shown in figure 4, most mutants retained between 40% and 90% of WT t^6^A activity *in vitro*. However, two mutants C110R^OSGEP^ and G177A^OSGEP^ involve residues located in the active site groove of OSGEP and exhibit significantly reduced t^6^A activity (4% and 15% of residual activity respectively). The highly conserved C110^OSGEP^ features a γS atom positioned 5.6 Å from the threonyl moiety of the TC-AMP analog, while G177^OSGEP^ interacts with the adenine of TC-AMP (Fig. S14A). Their respective mutations into arginine and alanine likely disrupt TC-AMP intermediate binding, explaining the substantial decrease in t^6^A activity. Additionally, the strictly conserved E151^OSGEP^ is part of the G150-D154 loop and forms a salt bridge with R325^OSGEP^ that may interact with the catalytically important C-terminus of TP53RK (Fig. S14B). E151^OSGEP^ and R325^OSGEP^ form a salt bridge positioned near the C-terminal helix of TP53RK, and the E151K^OSGEP^ mutation leads to a 32% residual activity. Additionally, three GAMOS mutations occur within R325^OSGEP^, resulting in t^6^A activities ranging from 40% to 75% of WT.

**Figure 3.**
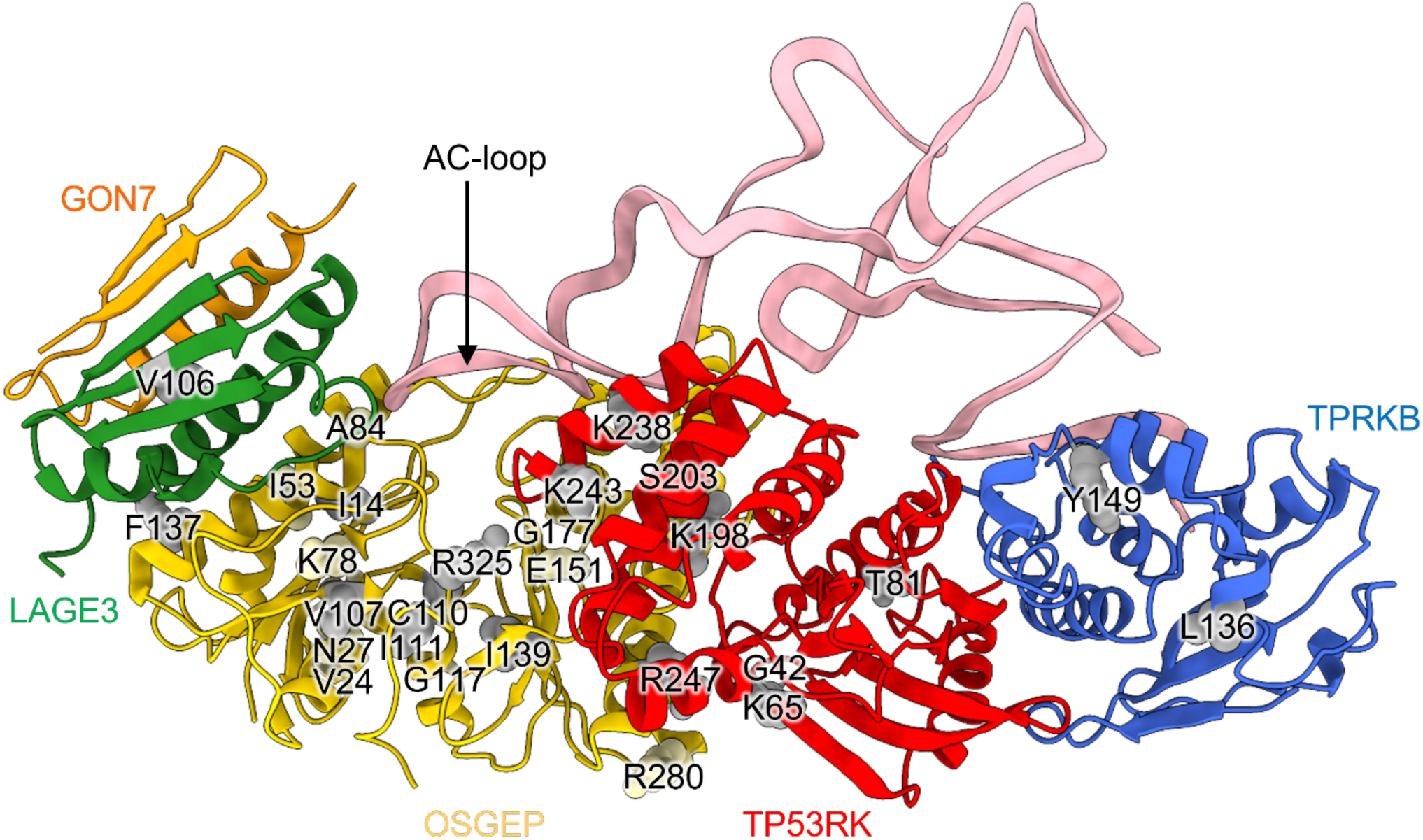
GAMOS mutations mapped onto the structure of the hKEOPS/tRNA. Overall view of the mutations. grey spheres: mutations studied *in vitro* (t^6^A and ATPase activity assays + tRNA binding) and *in vivo* in humanized yeast (Fitness) ; light-yellow spheres: mutations studied only *in vitro* (t^6^A and ATPase activity assays + tRNA binding). Colored subunits are orange for GON7, green for LAGE3, yellow for OSGEP, red for TP53RK, blue for TPRKB and pink for tRNA^Ile^_AAU_.

**Figure 4.**
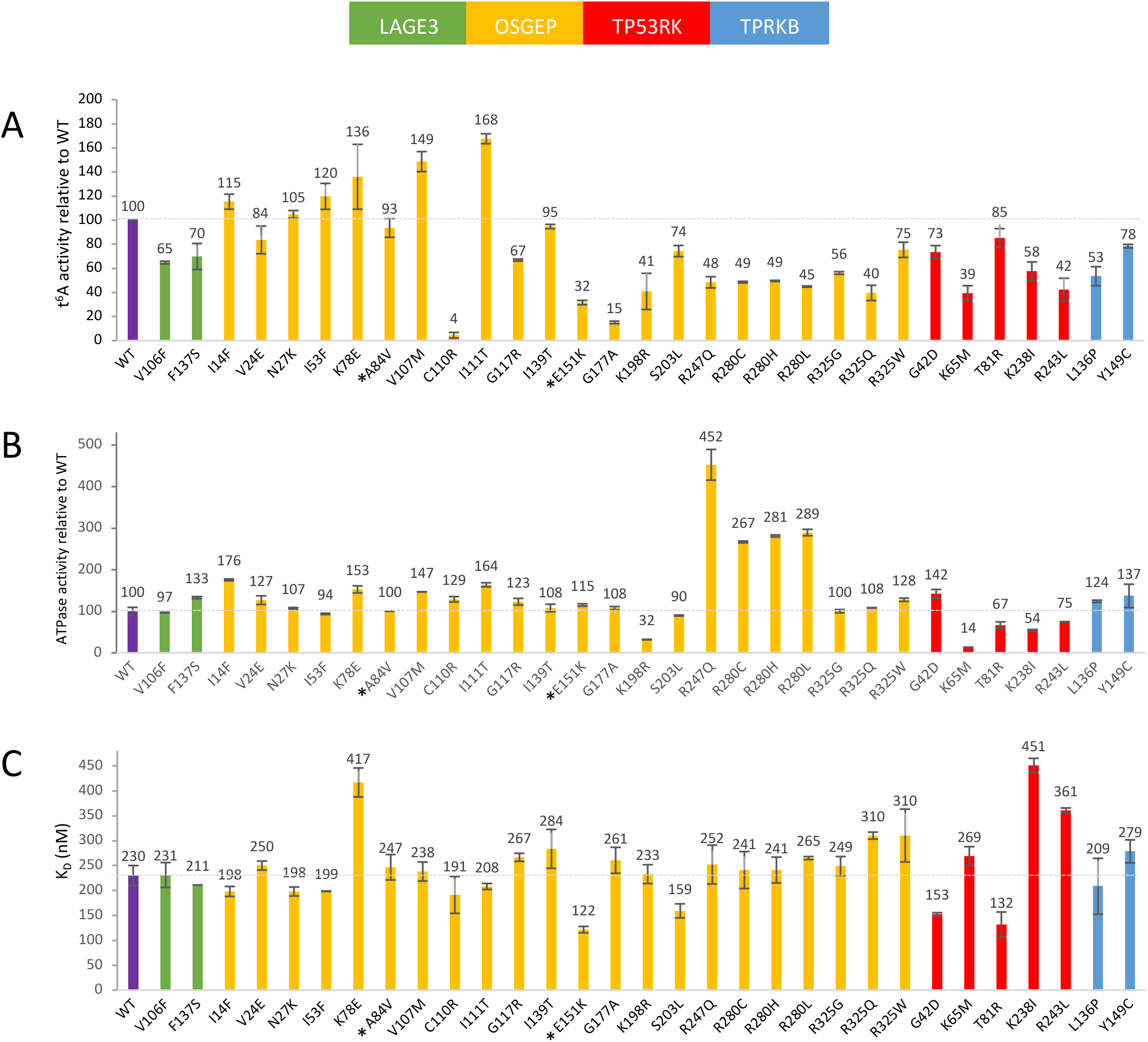
Biochemical characterization of Galloway-Mowat hKEOPS mutants. All experiments were performed using *in vitro* transcribed tRNA^Ile^_AAU_. t^6^A and ATPase activities are reported relative to WT hKEOPS (values shown as mean ± s.d.). (A) t^6^A modification activity, assessed by measuring the incorporation of radiolabeled threonine into tRNA. (B) tRNA-dependent ATPase activity, measured using an enzymatic ATP-regenerating assay (see supplementary methods). (C) Binding affinities of hKEOPS mutants for tRNA, determined by fluorescence anisotropy (see supplementary methods). * A84V + E151K : new heterozygous pathogenic variant described in table S22

Remarkably, two mutations that severely impair t^6^A activity C110R^OSGEP^ and E151K^OSGEP^ are consistently found as heterozygotes in the GAMOS population, associated with alleles causing a less drastic reduction in activity. Specifically, C110R^OSGEP^ is found in combination with I111T^OSGEP^, G177A^OSGEP^, R280C^OSGEP^, or R325Q^OSGEP^, while E151K^OSGEP^ is associated with A84V^OSGEP^ (Table 1). This pattern suggests that t^6^A levels below a critical threshold may be incompatible with cell viability, making the homozygous versions of C110R^OSGEP^ and E151K^OSGEP^ potentially lethal. Interestingly, five mutants exhibited significantly higher t^6^A activity than WT: I14F^OSGEP^, I53F^OSGEP^, K78E^OSGEP^, V107M^OSGEP^, and I111T^OSGEP^. This led us to question whether their Michaelis-Menten constant (K_M_) might be affected. Since our end-point assay does not provide kinetic parameters (k_cat_ and/or K_M_), we performed tRNA saturation experiments for WT hKEOPS and three of these mutants (I14F^OSGEP^, V107M^OSGEP^ and I111T^OSGEP^, as well as for a weakly active mutant (R325Q^OSGEP^) as a control (Fig. S15).

**Table 1.** *In vivo* and *in vitro* functional characterization of GAMOS-associated mutations in the t^6^A biosynthesis pathway. Individual GAMOS-associated mutations were introduced into the fully humanized yeast strain as described in Materials and Methods. The resulting mutant strains were evaluated for growth fitness and for the level of t⁶A modification (see supplementary methods) present in purified total tRNAs. In parallel, complete gene deletions were performed for each component of the human t⁶A biosynthesis pathway and similarly assessed. For mutations located in OSGEP, LAGE3, TP53RK, and TPRKB, full hKEOPS complexes harboring the respective mutant subunits were expressed in *E. coli*, purified and tested for *in vitro* enzymatic activity assays. As GAMOS is a recessive disorder, allele combinations identified in patients are also indicated (hom.: homozygous; het.: heterozygous, with the associated allele in parentheses; hemi.: hemizygous). nd: not detectable; n/a: not applicable; *: data taken from figure 4

| Yeast strain | t<sup>6</sup>A level<br>(% of wt) | Fitness | In vitro activities of<br>mutated hKEOPS (% of wt) |  | Allele combinations observed<br>in GAMOS patients |
| --- | --- | --- | --- | --- | --- |
|  |  |  | t<sup>6</sup>A synthesis* | ATPase* |  |
| IMX672 | 76 | + | / | / |  |
| humanized (wt) | 100 | + | 100 | 100 |  |
| *OSGEP* Δ | nd | - | n/a | n/a |  |
| I14F | 131 | + | 115 | 176 | hom. |
| V107M | 108 | + | 149 | 147 | hom. / het.(G177A) |
| C110R | nd | - | 4 | 129 | het.(I111T, G177A, R280C, R325Q) |
| I111T | 136 | + | 168 | 164 | het.(C110R) |
| I139T | 88 | + | 95 | 108 | hom. |
| G177A | 103 | + | 15 | 108 | hom. / het.(V107M, C110R) |
| K198R | 75 | + | 41 | 32 | het. (R247Q) |
| R247Q | 114 | + | 48 | 452 | hom. / het.(K198R) |
| R325Q | 51 | + | 40 | 108 | hom. / het.(C110R, R280H) |
| R325W | 19 | +/- | 75 | 128 | het.(K78E) |
| *LAGE3* Δ | 16 | - | n/a | n/a |  |
| F137S | 74 | + | 70 | 133 | hemi. |
| V106F | 90 | + | 65 | 97 | hemi. |
| *TP53RK* Δ | nd | - | n/a | n/a |  |
| G42D | 85 | + | 73 | 142 | hom. |
| K65M | 37 | + | 39 | 14 | hom. |
| T81R | 81 | + | 85 | 67 | het.(K60Sfs ) |
| K238I | 71 | + | 58 | 54 | hom. |
| R243L | 67 | + | 42 | 75 | hom. |
| D162A | 84 | + | 64 | 0 |  |
| *TPRKB* Δ | nd | - | n/a | n/a |  |
| L136P | 77 | + | 53 | 124 | hom. |
| Y149C | 78 | + | 78 | 137 | hom. |
| *GON7* Δ | 77 | + | n/a | n/a |  |
| Y7stop | 83 | + | n/a | n/a | hom. |
| Y7Lfs | 85 | + | n/a | n/a | hom. |
| *YRDC* Δ | nd | - | n/a | n/a |  |
| A84V | 87 | + | n/a | n/a | het.(V241lfs ) |
| V241lfs | nd | - | n/a | n/a | het.(A84V) |
| L265Δ | 85 | + | n/a | n/a | hom. |

The V_max_ (or k_cat_) values for the three active mutants were higher *in vitro* than those of the WT. However, their K_M_ values were also increased, leading to an overall decrease in their *in vitro* catalytic efficiency (k_cat_/K_M_). This suggests that most GAMOS mutations may impact the overall activity of hKEOPS, either by reducing the catalytic constant (k_cat_) and/or increasing the K_M_ value. As a result, these mutations could lead to suboptimal levels of t^6^A-modified tRNAs in human cells under physiological conditions. Twelve OSGEP mutations are found as six pairs of neighboring residues surrounding the active site groove. Four of these pairs: V24E^OSGEP^/N27K^OSGEP^, I14F^OSGEP^/I53F^OSGEP^, K78E^OSGEP^/V107M^OSGEP^, and I111T^OSGEP^/I139T^OSGEP^ harbor GAMOS mutations with near-WT or even higher t^6^A activities. In contrast, two pairs, E151K^OSGEP^/R325G^OSGEP^ and K198R^OSGEP^/S203L^OSGEP^ exhibit reduced t^6^A activities.

K198^OSGEP^ is within hydrogen-bonding distance of the main chain of G161^TP53RK^, which neighbors the catalytic residue D162^TP53RK^ (Fig. S14C). The K198R^OSGEP^ mutation weakens both t^6^A (41%) and ATPase (30%) activities. R247^OSGEP^ does not directly interact with TP53RK in the cryo-EM structure but forms a salt bridge with D162^TP53RK^ in the AF3 model (Fig. S14D). Interestingly, the equivalent residue in archaeal KEOPS has been proposed to act as a communicator between t^6^A and ATPase activities. In archaeal KEOPS, mutating this residue to alanine completely abolished both activities, suggesting a role in stabilizing the γ-phosphate of ATP during hydrolysis. However, our biochemical data do not support this hypothesis. In fact, R247Q^OSGEP^ retains 50% of its t^6^A activity while exhibiting a 4.5-fold increase in ATPase activity. Similarly, three GAMOS mutations at residue R280^OSGEP^ (R280C^OSGEP^, R280H^OSGEP^ and R280L^OSGEP^) retain about 50% t^6^A activity but show a striking increase in ATPase activity (∼280%). While R280^OSGEP^ does not interact with TP53RK in the cryo-EM structure (Fig. S14B), it forms a hydrogen bond with the main-chain carbonyl of G42^TP53RK^ in the P-loop in the AF3 model (Fig. S14D). The behavior of these mutants highlights the complex and poorly understood relationship between the ATPase and t^6^A activities of KEOPS.

In the AF3 model, A84^OSGEP^ contacts the tRNA A_34_ base (Fig. S14E), but the A84V^OSGEP^ mutation has little impact on t^6^A activity or tRNA affinity. G117^OSGEP^ is surrounded by hydrophobic side chains (Fig. S14F), and the G117R^OSGEP^ mutation results in a 30% loss of t^6^A activity. While we initially expected this mutant to be unstable, no such instability was observed during purification. In the AF3 model, S203^OSGEP^ is near the C_10_ ribose of tRNA that has been shown to be important for the recognition of cognate tRNAs by the hKEOPS machinery (Wang et al., 2022). The S203L^OSGEP^ mutation reduces t^6^A activity by 25% but increases tRNA affinity.

All GAMOS mutations within the TP53RK subunit result in weakened t^6^A activity, ranging from 39% to 85% of WT. G42^TP53RK^ is a fully conserved residue within the P-loop (Fig. S16A), and the G42D^TP53RK^ mutation retains 75% of t^6^A activity while increasing ATPase activity to 140%. In the AF3 model, K65^TP53RK^ interacts with the main-chain carbonyl of A43^TP53RK^ in the P-loop (Fig. S16B). The K65M^TP53RK^ mutation reduces ATPase activity (15% of residual activity) but maintains 39% of WT t^6^A activity.

The biochemical activities of T81R^TP53RK^ remain largely unchanged, which aligns with its positioning, neither near the ATPase binding site nor close to tRNA (Fig. S16A-B). In contrast, K238^TP53RK^ and R243^TP53RK^, located in the C-terminal α-helix that interacts with the tRNA anticodon stem (Fig. S16C), show significantly reduced tRNA affinity. Mutations K238I^TP53RK^ and R243L^TP53RK^ lead to a 50–60% loss of t^6^A activity, further emphasizing the crucial role of TP53RK’s C-terminal helix in t^6^A modification, as previously suggested by site-directed mutagenesis studies (this paper and (Ona Chuquimarca et al., 2024)).

The two GAMOS mutations in TPRKB (L136P^TP53RK^ and Y149C^TP53RK^) involve residues that do not directly contact tRNA (Fig. S16D) but still lead to reduced t^6^A activity. Although L136^TP53RK^ and Y149^TP53RK^ are embedded in a hydrophobic environment, their mutations to proline and cysteine, respectively, do not compromise hKEOPS stability.

Similarly, V106^LAGE3^ is part of the hydrophobic core of LAGE3, and its mutation to phenylalanine causes a 35% reduction in t^6^A activity without affecting stability or tRNA affinity. F137^LAGE3^, which interacts with a hydrophobic patch on OSGEP, is far from OSGEP’s active site pocket, yet its mutation to serine results in a 30% loss of t^6^A activity (Fig. S16E).

### 4. Evaluation of GAMOS-associated alleles in cellulo

Intrigued by the observation that most GAMOS mutations retained over 50% of WT t^6^A activity *in vitro*, we wondered whether these t^6^A levels would also be maintained in a cellular context. Since obtaining biological material from rare disease patients for biochemical analysis is nearly impossible, we opted for a reverse genetics approach to investigate the impact of GAMOS mutations on cell fitness and t^6^A levels *in vivo*. Yeast was our model of choice due to its highly similar t^6^A system to humans and its ease of genetic manipulation. To make this approach feasible, we used a haploid yeast strain in which the entire endogenous t^6^A pathway was replaced with its human orthologues using CRISPR-Cas9. An explanation of this strategy can be found in the supplementary methods.

The yeast strain fully humanized for the t^6^A enzymatic pathway remained viable, with t^6^A levels comparable to those of the original strain (Fig. 5, Table 1). This suggests that the human t^6^A machinery functions effectively in yeast, successfully modifying yeast tRNAs. Additionally, the introduced human genes, except for *hGON7*, became essential for yeast cell fitness, as their individual deletion caused significant growth defects and a sharp decline in t^6^A levels (Fig. 5B). In contrast to its yeast homolog *yGON7*, deleting *hGON7* in the humanized yeast strain had no impact on cell fitness, confirming its less-essential role, consistent with previous knockdown experiments in human podocytes (Arrondel et al., 2019).

**Figure 5.**
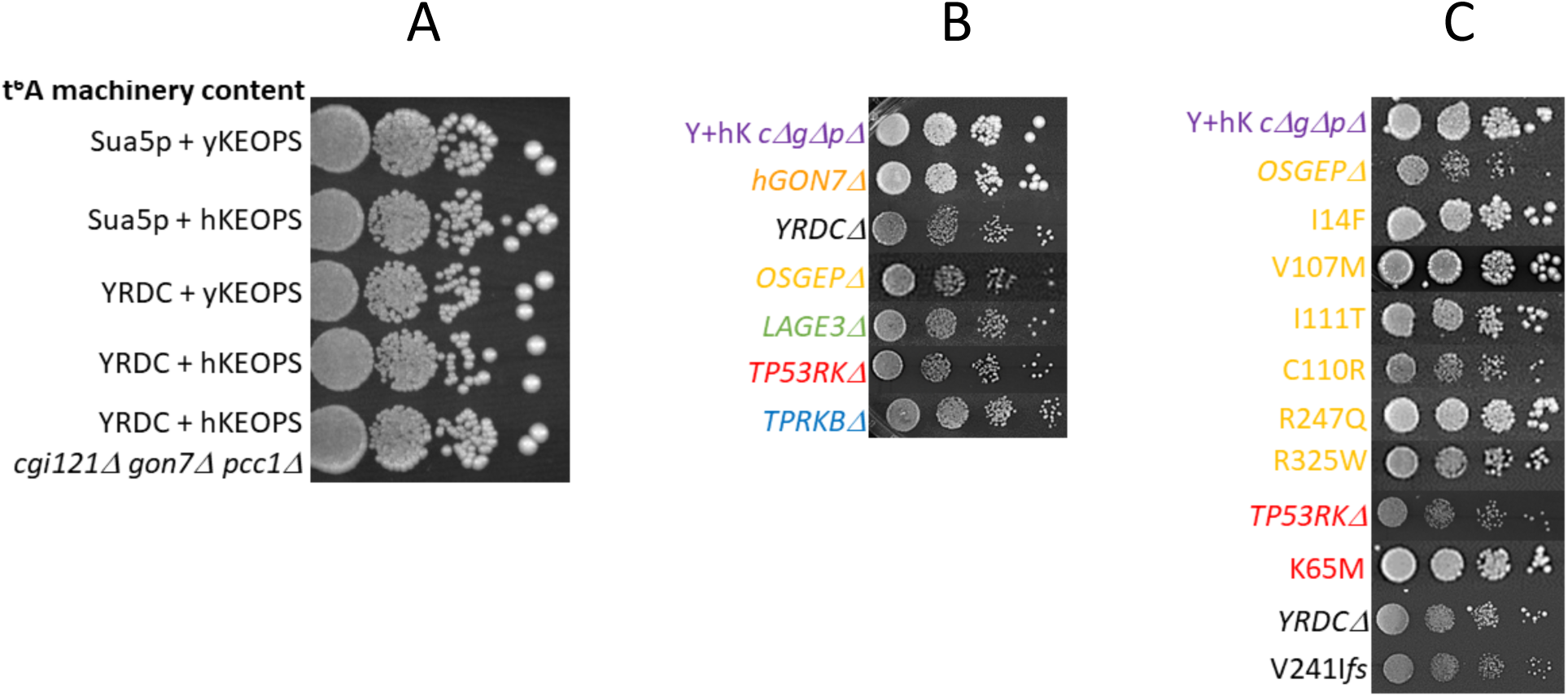
Yeast strain fitness assays. Plasmid-cured yeast strain cells were resuspended in water and subjected to 10-fold serial dilutions. 10 µL of each suspension were spotted onto agar plates and allowed to dry. Plates were incubated at 28 °C for 5 days before observation. (A) Replacement of the yeast t⁶A pathway proteins (ySua5 + yKEOPS) with their human homologs (YRDC + hKEOPS) results in a normal growth phenotype. Comparable growth is observed in yeast/human hybrid combinations (ySua5 + hKEOPS and YRDC + yKEOPS). The bottom row shows the fully humanized strain, in which the three vestigial yeast genes of the t⁶A pathway (*CGI121*, *yGON7*, *PCC1*) have been further deleted (YRDC + hKEOPS *cgi121Δ gon7Δ pcc1Δ*). This strain, referred to as Y+hK *cΔgΔpΔ* in the following panels, was used as the recipient strain for all CRISPR/Cas9 mutagenesis experiments. (B) Deletion of individual genes from the human t⁶A pathway reduces the fitness of the fully humanized yeast strain, except in the case of *hGON7*. The top row shows the growth of the intact humanized strain (Y+hK *cΔgΔpΔ*). (C) The impact of selected GAMOS-associated mutations in OSGEP, TP53RK, and YRDC on the fitness of the fully humanized yeast strain was assessed. Growth of the unmutated humanized strain (Y+hK *cΔgΔpΔ*) serves as the normal growth control. For comparison, the effects of individual deletions of *OSGEP*, *TP53RK*, and *YRDC* are also shown. Among the GAMOS mutants, only C110R^OSGEP^, R325W^OSGEP^ (to a lesser extent), and V241I*fs*^YRDC^ show impaired growth.

Among the 25 GAMOS mutations introduced individually into the humanized yeast strain, only two (V241I*fs*^YRDC^ and C110R^OSGEP^), severely impaired cell fitness, with effects comparable to a full gene deletion (Fig. S17 and Table 1). The R325W^OSGEP^ mutation caused a slight fitness reduction, while the remaining 21 mutations exhibited normal growth. We supplemented these fitness assessments by measuring t^6^A levels in tRNAs isolated from growing cells. For V241I*fs*^YRDC^ and C110R^OSGEP^, no detectable t^6^A was observed, mirroring the results for *YRDCΔ* and *OSGEPΔ*. In contrast, R325W^OSGEP^ retained 19% of the WT t^6^A level, while the other mutants ranged from 37% (K65M^TP53RK^) to 136% (I111T^OSGEP^). Since R325W^OSGEP^ reduced fitness while K65M^TP53RK^ did not, we estimate that the t^6^A threshold for normal growth in yeast lies between 19% and 37% of the WT level. Four OSGEP mutants (I14F^OSGEP^, V107M^OSGEP^, I111T^OSGEP^, R247Q^OSGEP^) displayed elevated *in vivo* t^6^A levels. For three of these (I14F^OSGEP^, V107M^OSGEP^, I111T^OSGEP^), the increase aligned with their hyperactivity in both t^6^A synthesis and ATP hydrolysis observed *in vitro*, while R247Q^OSGEP^ exhibited a depressed level in t^6^A synthesis and an ATPase activity boosted 4.5 times compared to WT (Table 1).

Clinically, the GAMOS alleles that most severely impair t^6^A activity (V241I*fs*^YRDC^, C110R^OSGEP^, R325W^OSGEP^) consistently appear in heterozygous combinations with alleles supporting normal growth and maintaining at least 51% of the WT t^6^A level (e.g., R325Q^OSGEP^, see Table 1). In these cases, the second allele may partially compensate for the deleterious mutation, potentially supporting fetal development and embryonic viability. Although we have not tested the K60S*fs*^TP53RK^ allele—anticipating that its phenotype would resemble that of a full deletion—the aforementioned pattern appears to hold: this allele has only been observed in patients as a heterozygous combination with the T81R^TP53RK^ variant, which maintains a sufficient level of t⁶A. Notably, all GAMOS-associated alleles tested in this study and identified in patients in the homozygous state (or hemizygous for X-linked LAGE3) maintain at least 37% of WT t⁶A levels (e.g., K65M^TP53RK^).

Two homozygous nonsense GAMOS mutations were identified at position 7 in *hGON7*, one introducing a stop codon and another leading to a frameshift. Both mutations were previously shown to cause the absence of protein expression. We tested the performance of these mutations in yeast humanized for KEOPS. The two point-mutants, as well as the complete deletion of the *hGON7* gene, were fully viable and produced near WT levels of t⁶A, confirming that *hGON7* is less essential than the other subunits of the complex. Nevertheless, its absence in human cells may still contribute to a milder form of the disease.

We finally tested the properties of the ATPase catalytic (non-GAMOS) mutant, D162A^TP53RK^. Mutation of the equivalent residue in archaeal KEOPS annihilates both the ATPase and t^6^A activities (Ona Chuquimarca et al., 2024). D162A^TP53RK^ has no ATPase activity, but kept 64% of its t^6^A activity *in vitro,* showing that the ATPase activity is not absolutely required *in vitro* for t^6^A synthesis in hKEOPS. Intrigued by this surprising result, we introduced this mutation into yeast, humanized for KEOPS. The D162A^TP53RK^ mutant is perfectly viable and t^6^A yields are 84% of WT levels, in line with our *in vitro* data. We further showed that the ATPase activity of hKEOPS requires the presence of cognate tRNA (Fig. S18). We conclude that the necessity of ATP hydrolysis very much depends upon organism and also explains why the R247Q^OSGEP^ keeps 50% of its t^6^A activity, while the alanine mutation of the equivalent residue in archaeal KEOPS is inactive.

## Discussion

The cryo-EM structure of tRNA-bound hKEOPS shows that hKEOPS maintains a linear configuration establishing contacts for 4 out of 5 subunits with the L-shaped tRNA. The tRNA binding mode brings its AC-loop near the active site groove of OSGEP and its CCA tail is bound to TPRKB. The affinity is explained by a good complementarity between tRNA and the positively charged electrostatic surface patches of hKEOPS. Very few base-specific interactions occur between hKEOPS and tRNA, in line with the comparable binding affinities that we measured with substrate and non-substrate tRNAs. However, the strategy to obtain good shape complementarity is somewhat different between human and archaeal complexes. The structures of KEOPS in the apo- and tRNA bound forms are superposable in the case of archaeal but not for human KEOPS. Flexibility of the OSGEP/TP53RK interface, enables the hKEOPS complex to embrace the 3D shape of the substrate tRNA. The only direct interaction of hKEOPS with bases from the tRNA is between TPRKB and the CCA tail, confirming the CCA interaction with Cgi121 of the archaeal complex, although the orientation of the tails is different in the two complexes.

The AC-loop, although poorly defined in the hKEOPS structure, obviously does not penetrate the active site pocket. The cryo-EM data collected on an archaeal KEOPS/tRNA complex revealed the presence of at least two tRNA conformations. One of those was similar to the tRNA conformation in the hKEOPS complex, while the other was a distorted conformation with a presumably released A_37_ base (Ona Chuquimarca et al., 2024). Although the lower resolution of the tRNA region in our maps suggests mobility, we did not find evidence for the latter tRNA conformation in the human complex. Interestingly, a cryo-EM structure of the tRNA deaminase ADAT2/3 reveals a more open conformation of the AC-loop, exposing bases 34 and 37 of the substrate tRNA (Dolce et al., 2022) (Fig. S19). Although the modification in the latter case occurs at position 34, this observation supports the notion that the tRNA anticodon loop might be intrinsically flexible and could adopt distinct conformations along the modification reaction. The AF3 model of the hKEOPS/tRNA complex suggests how a simple rearrangement of the AC backbone combined with the pivoting of A_37_ (and also A_34_) could bring its N^6^ amino group in front of the carbamoyl moiety of TC-AMP. In this model, the loop contained between residues (S132-N135)^OSGEP^ is well positioned to stabilize the oxyanion generated during the reaction (distance between carbamoyl oxygen of TC-AMP and main chain nitrogens of G133 and G134: 3.5 Å; mutant S132A^OSGEP^ causes 50 % loss of t^6^A activity) (Fig. S20). We therefore presume that in absence of the TC-AMP intermediate, tRNA is bound in a pre-catalytic conformation. This binding mode could correspond to those of non-cognate tRNAs. We speculate that the binding of the TC-AMP intermediate triggers conformational changes in hKEOPS and tRNA that would promote the pivoting of A_37_ into the active site pocket, bringing its N^6^-aminogroup close to the carbamoyl moiety. An alternative, but not exclusive recognition mechanism, could be governed by the ATPase activity of TP53RK. It was indeed observed for archaeal KEOPS that a strong coupling exists between t^6^A and ATPase activities, but the t^6^A activity of hKEOPS does not require an active ATPase. Clearly the relationship between both activities is complex and dependent upon organism. This complexity is further emphasized by the fact that the mitochondrial Kae1 paralog is able to synthesize t^6^A on its own.

The cryo-EM structures of the archaeal and hKEOPS complexes bound to tRNA, likely provide different snapshots of the t^6^A reaction cycle. The latter could start by the binding of a tRNA CCA tail to TPRKB followed by a conformational change of the hKEOPS to bring the active site of OSGEP near the AC-loop. The release of the A_37_ base out of the core of the AC-loop could be triggered by a conformational change of the tRNA and/or by the binding of the TC-AMP intermediate. It was shown for the archaeal KEOPS/tRNA complex that t^6^A activity required substrate tRNA induced activation of the Bud32 ATPase. This mechanism nicely explains how, in absence of base specific interactions, KEOPS could distinguish between cognate and non-cognate tRNAs. However, the ATPase inactive D162A^TP53RK^ mutant, keeps 70% of its t^6^A activity *in vitro*. Extrapolating from archaeal mutant studies, it was proposed that R247^OSGEP^ could activate the ATPase activity of TP53RK by binding to the γ-phosphate of ATP. The GAMOS mutant R247Q^OSGEP^ mutant however has a 4-fold higher ATPase and 50% t^6^A activity. These observations highlight that the very similar structures of the human and archaeal KEOPS complexes may carry significant mechanistic differences.

Although rare diseases are genetically diverse, their biochemical effects often converge on a few major pathways: loss of function, toxic gain of function, metabolic disruption or protein misfolding. GAMOS is characterized by a large number of mutations in different proteins. Six of these proteins (YRDC and hKEOPS subunits) constitute the biosynthesis pathway of the ancient and essential t^6^A tRNA modification. It was therefore expected that the GAMOS mutations would invariably lead to the loss of this function. However, a more complex picture emerges from a detailed analysis of the behavior of these GAMOS mutations. First, our biochemical data suggest that these mutations do not suffer from protein misfolding and that they do not affect the stability of the quaternary structure of hKEOPS. Our structural data are in line with this conclusion. Mapping of the GAMOS mutations onto the structure of the hKEOPS/tRNA complex indeed shows that they are (1) positioned at the protein surface and hence should not affect protein stability and (2) rarely involved in subunit interactions. Secondly, apart from C110R^OSGEP^ and G177A^OSGEP^, all the mutations have moderate (above 30%) to high t^6^A activities *in vitro*. These activities are for most of the mutants in good agreement with the t^6^A levels measured from the mutated yeast strains. Also, the fitness of most of the mutants was undistinguishable from that of the WT hKEOPS strain. The fitness of the R325W^OSGEP^ mutant (19% t^6^A level) was reduced, whereas that of the K65M^TP53RK^ mutant (37% t^6^A level) was not. This suggests that, in our yeast model system, a t^6^A level below 20% may be deleterious for cell viability.

In human GAMOS pathology, we observed that highly severe mutations (i.e., C110R^OSGEP^ or E151K^OSGEP^) are mostly associated with a milder mutation in heterozygous individuals and are never found in the homozygous state. We therefore presume that, in humans, a severe impairment of the t^6^A pathway may not be compatible with fetal development or embryonic viability. However, a reduced but not completely absent cellular t^6^A level may lead to the pathology.

Several neurodevelopmental disorders appear to be linked to defects in tRNA modifications. It has been established that neurons and podocytes are particularly sensitive to disturbances in their tRNA modification pathways (Braun et al., 2018b; Chujo and Tomizawa, 2025; Suzuki, 2021; Zhang and Westhof, 2025). Overall, our observations on GAMOS mutations argue against a simple loss-of-function mechanism of the t^6^A machinery. Instead, they support a model in which these mutations impair KEOPS catalytic efficiency below a critical threshold required for optimal cellular homeostasis

## MATERIALS AND METHODS

Strains, media and reagents are mentioned in supplementary methods.

### hKEOPS/tRNA sample preparation

Directly after purification, hKEOPS was mixed with human tRNA^Ile^_AAU_ (previously stored at - 20°C) in a molar ratio of 1:1.3. To optimize the interaction the buffer contained a final concentration of 50 mM NaCl. After 30 min incubation at 18°C, the sample was purified by size exclusion chromatography using a Superdex^TM^ 200 Increase 10/300 GL column (Cytiva). The buffer used in the size exclusion column consisted of 20 mM HEPES pH 7.5, 50 mM NaCl, 5 mM 2-mercaptoethanol. Fractions on top of the peak containing the hKEOPS-tRNA complex were isolated and stored overnight at 4°C. The sample was used directly the following day for cryo-EM experiments.

### Grid preparation and cryo-EM data collection

Negatively stained electron microscopy images showed good homogeneity and distribution of the complex. After optimizing the cryo-grid preparations, R2/2 holey-carbon coated gold 300 mesh grids (Quantifoil^®^) were glow discharged twice for 25 s at 15 mA before sample freezing. Three microliters of sample at 0.35 mg/mL concentration were placed on the grid, blotted for 8.0 s, and flash-frozen in liquid ethane using a Vitrobot Mark IV (Thermo Scientific^TM^) operated at 4°C and 100% humidity. The electron microscopy data collection statistics are gathered in figure S2. A total of 14328 micrographs were collected in a EER format, on a Krios G4i microscope at the NanoImaging Core Facility at Institut Pasteur (Paris) operating at 300 kV equipped with a Falcon 4i direct electron detector and a Selectris X imaging filter (Thermo Scientific^TM^). The automation of the data collection was done with the software EPU. Movies were recorded at 165000x nominal magnification in electron-counting mode using exposures of 2.16 s with a dose rate of 18.48 electrons/Å^2^/s, resulting in a total dose 40 electrons/Å^2^. 14328 movies were acquired with a defocus range from −3.0 to −1.0 μm. The pixel size was 0.77 Å per pixel.

### Cryo-EM data processing

Data were processed using CryoSPARC v4.2.1 (Punjani et al., 2017). The movies were motion-corrected using patch motion correction and contrast transfer function estimation was done using CTFFIND4 (Rohou and Grigorieff, 2015). Micrographs were screened using curate exposure according to CTF fit resolution (at least 6.5 Å) and total full-frame motion distance (less than 20 Å). 2D templates for picking were obtained by a first automatic (blob) picking over the 13411 previously selected micrographs and followed by 2D classifications. 14 good 2D class averages were used for template picking with a particle diameter of 140 Å. A total of 9613829 particles were extracted using a box size of 246 Å, with a down-sampling factor of 4. To clean up the particle set, two successive iterations of 2D classifications were performed. For each iteration, the particles were sorted into 200 different classes, and a selection of the best classes was made. This process produced two different selections. A first selection of 34 2D class averages corresponding to 552340 particles for an object close to the expected size, and a second selection of 36 2D class averages corresponding to 315314 particles for an object smaller than expected. Reconstructions of an initial three-dimensional map model from our particle selections were performed using ab-initio reconstruction. We obtained 4 distinct volumes of varying size. Particle selection of a small-sized object enabled the calculation of a low-resolution volume of a single tRNA. While analysis for the larger object yielded 3 distinct low-resolution volumes corresponding to i) hKEOPS in complex with tRNA, ii) apo-hKEOPS, and iii) a tRNA interacting with an object smaller than full hKEOPS which we interpreted as probably a sub-complex. Refinement of the three-dimensional map was not carried out for tRNA alone. Refinement of the other three maps was carried out using heterogeneous refinement, which refines the various initial models while sorting the particles between the different volumes (here three). After particle re-extraction with a box size of 246 Å again, but without down-sampling, further refinement of the three different volumes was carried out independently. 214527 particles were used for the volume corresponding to apo-hKEOPS, 214607 for the hKEOPS/tRNA complex and 121547 for the hKEOPS/tRNA sub-complex. All three maps were refined using Non-Uniform Refinement (Punjani et al., 2020), which provided the best results here. The two maps corresponding respectively to apo-hKEOPS and to the hKEOPS/tRNA subcomplex were also refined with two additional processing steps: a Global CTF refinement step followed by a second Non-Uniform Refinement. The 3D map KEOPS/tRNA showed no improvement in quality or resolution after these two additional steps. Gold Standard Fourier Shell Correlation was used to determine the resolution of the final maps: 3.9 Å for the hKEOPS/tRNA, complex 3.7 Å for apo-hKEOPS, and 4.2 Å for the hKEOPS/tRNA sub-complex. Figure S21 shows the complete cryo-EM processing workflow.

### Model building and refinement

The building of initial models into the experimental density maps was guided by the available crystal structures of the sub-complexes OSGEP-LAGE3-GON7 (PDB ID, 6GWJ) (Arrondel et al., 2019) and TPRKB-TP53RK (PDB ID, 6WQX) (Li et al., 2021). For the tRNA, the crystal structure of *E. coli* aspartate tRNA^Asp^_UGU_ (PDB ID, 6UGG) (Chan et al., 2020) have been used due to similar secondary structures and nucleotides have been mutated with COOT (Emsley et al., 2010) to match the sequence of the human tRNA^Ile^_AAU_ used in the experiments. The atomic models were placed into the maps by rigid body fitting using either the Dock in Map tool available in PHENIX (Liebschner et al., 2019) or the fit in map tool available in Chimera (Pettersen et al., 2004). Several iterative cycles of flexible fitting using the Namdinator server (Kidmose et al., 2019) with a gradual increase in G-force parameter coupled with refinement using the real space refinement tool in PHENIX and manual refinement in COOT provided the final cryo-EM atomic models. Structure figures were generated using Chimera, ChimeraX (Meng et al., 2023) and PyMOL (The PyMOL Molecular Graphics System, Version 2.0 Schrödinger, LLC) software. Refinement statistics are summarized in figure S2D.

### *In vitro* t^6^A activity assays

The t^6^A modification enzyme activity assay was based on a stopped assay monitoring the incorporation of radioactive ^14^C-Threonine into the tRNA substrate (see supplementary materials) after 10 minutes incubation at 30°C (Daugeron et al., 2023; Perrochia et al., 2013a). It further requires the use of a coupled enzyme system: (YrdC or ySua5) to supply the ^14^C-TC-AMP intermediate and pyruvate kinase + PEP to regenerate the ATP consumed by the ATPase activity of the TP53RK subunit. In a final volume of 25 µL, human tRNA^Ile^_AAU_ (14 µM) was incubated with 150 µM L-threonine [1-^14^C] (55 mCi/mmol, Isobio), 50 mM Hepes pH 8, 35 mM KCl, 3.3 mM MgCl_2_, 10 µM ZnCl_2_, 5 mM DTT, 1 mM phosphoenolpyruvate, 0.5 mM ATP, 10 U/mL Pyruvate kinase, 5 µM ySua5 and 3 μM hKEOPS for 10 min at 32°C. 1 mL of 15% (w/v) trichloroacetic acid (TCA) was added to stop the reaction and the mixture was then incubated on ice for 1 h before loading on a pre-wet glass microfiber filter GF/F (Cytiva/Whatman) using a vacuum filtration apparatus. Filters were washed three times with 1 mL of 5% TCA and 1 mL of 95% EtOH, precooled to 4°C, and dried. Liquid scintillation counter (Hidex 300 SL) was used to quantify the t^6^A amount in tRNA.

### Introducing the GAMOS mutations in the humanized yeast strain

For each mutation a sgRNA-encoding DNA fragment and a repair fragment are required. The 20 bp sequence target has to be located the closest to the GAMOS mutation site and the repair fragment is designed to introduce the mutation but also to modify the restriction map of the repair fragment toward a given enzyme for restriction fragment length polymorphism (RFLP) diagnosis purpose (either creating or destroying a cutting site). To avoid multiple cycles of cut/repair in the cell, silent mutations were introduced in the repair DNA to destroy the 20bp target sequence and/or the NGG palindrome adjacent motif. In the case gaps existed between the target sequence, the GAMOS mutation and the modified restriction site, silent mutations were introduced within these gaps to encourage HR events outside of this region (*ie.* upstream and downstream) therefore preserving the link between the GAMOS mutation and the modified restriction site. The recipient strain for CRISPR/Cas9-mediated mutagenesis was the fully humanized yeast strain (Y+hK *cΔgΔpΔ ;* see supplementary materials) already transformed with a *TRP1* plasmid coding either for ySua5 (to preserve cell viability during *YRDC* mutagenesis) or for ΔN33Qri7p (to preserve cell viability during mutagenesis targeting any of the hKEOPS genes). The pMEL10 linear backbone (*URA3*) was systematically used. Triple transformations were performed as already described and selections were made on Synthetic Complete Medium lacking Trp and Ura. Genomic DNA was extracted from selected clones using the method published online (https://openwetware.org/wiki/Silver:_Colony_PCR). A 25-µL PCR reaction was run for each sample using HF-Phusion Polymerase (5*μ*L 5xHF Buffer, 0,5*μ*L dNTPs (10mM), 0.125*μ*L Forward Primer (100 *μ*M), 0.125*μ*L Reverse Primer (100 *μ*M), 0.25*μ*L de HF-Phusion Polymerase, 2.5*μ*L diluted Genomic DNA, 16.5*μ*L sterile H_2_O. Program: 30s at 98°C, 30s at the lowest melting temperature of the 2 primers, 15s at 72°C (repeated for 40 cycles). The PCR products were then purified on silica minicolumns using the GeneJet PCR Purification Kit from ThermoFisher and eluted in 25µL of elution buffer. For each sample a digestion using the appropriate restriction enzyme was performed in a final volume of 15µL (1.5µL 10x FastDigest Green Buffer; 1µL FastDigest Restriction Enzyme; 12.5µL Purified PCR Product) for 1h at 37°C. Samples were then loaded onto and run in a 2% agarose gel for RFLP. Positives clones were cured for their resident plasmids and finally the mutated region confirmed by sequencing.

### Evaluation of GAMOS associated alleles in human t^6^A pathway (Fitness assay)

Isolated clones from the plasmid-cured strains were resuspended in distilled water and cell density was adjusted to an OD at 600nm of 0.2 by dilution. Starting from this suspension, a 10-fold serial dilution was performed and 10µL of each dilution were spotted on SC medium. Plates were incubated for 5 days at 28°C prior observation.

### Data availability

Atomic coordinates of the apo-hKEOPS, hKEOPS/tRNA^Ile^_AAU_ complex and hKEOPS/tRNA^Ile^_AAU_ subcomplex have been deposited in the Protein Data Bank (PDB) with the ID codes 9FL9, 9T2U and 9T3H, respectively and the corresponding cryo-EM density maps in the Electron Microscopy Data Bank (EMDB) with the ID codes EMD-50536, EMD-55477 and EMD-55495, respectively.

## Acknowledgements

This work was supported by the Fondation pour la Recherche Médicale (project DEQ2015031682) (to C.Antignac), the European Union’s Seventh Framework Programme (FP7/2012, grant 305608 EURenOmics) (to C.Antignac), the Investments for the Future Program (grant ANR-10-IAHY-01) (to C.Antignac), ANR KeoGamo (ANR-18-CE11-0008-01) (to G.M., H.v.T. and C.V.B.). This work was supported by the French Infrastructure for Integrated Structural Biology (FRISBI) (ANR-10-INSB-05-01) (to H.v.T.). S.M. was supported by a PhD grant of the Fondation pour la Recherche Médicale (FRM).

Preliminary sample testing conditions have been obtained thanks to Instruct ERIC research infrastructure (INEXT Discovery program PID 5037) at the Cryo-EM facility of CSIC/CNB in Madrid, Spain.

The authors acknowledge the financial support from the Graduate School Life Sciences and Health (GS LSH) of Université Paris-Saclay, under AAP-Scientific-2021 grant (MOD’N’ART project) (to H.v.T. and D. T.). C.A.H.F. has received funding from the European Union Horizon Europe Research and Innovation Program under grant agreement no. 101026386. We thank the cryo-electron microscopy facility of I2BC and S. Bressanelli for their help. We thank the Nanoimaging Core facility at the Institut Pasteur and the help of J.-M. Winter, E. Salazar, S. Tachon, and M. Vos.

