## supplemental data for "Structure of the human KEOPS/tRNA complex and characterization of pathogenic variants responsible for the Galloway Mowat syndrome"

### SUPPLEMENTAL FIGURES

A

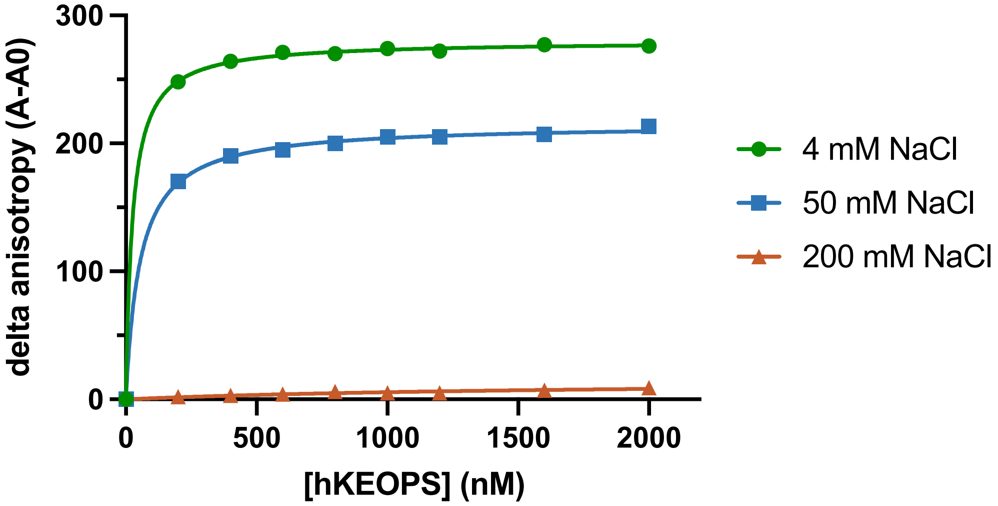


B

| Nonlin fit<br>Table of results | A<br>4 mM NaCl<br>Y | B<br>50 mM NaCl<br>Y | C<br>200 mM NaCl<br>Y |
|---|---|---|---|
| **One site -- Specific binding** | | | |
| **Best-fit values** | | | |
| Bmax | 280.1 | 215.2 | 15.09 |
| Kd | 25.29 | 54.44 | 1694 |
| **Std. Error** | | | |
| Bmax | 0.9020 | 1.197 | 4.252 |
| Kd | 1.707 | 3.381 | 824.1 |
| **95% CI (asymptotic)** | | | |
| Bmax | 277.9 to 282.2 | 212.4 to 218.0 | 5.032 to 25.14 |
| Kd | 21.25 to 29.32 | 46.45 to 62.44 | -254.7 to 3643 |
| **Goodness of Fit** | | | |
| Degrees of Freedom | 7 | 7 | 7 |
| R squared | 0.9998 | 0.9993 | 0.9267 |

C

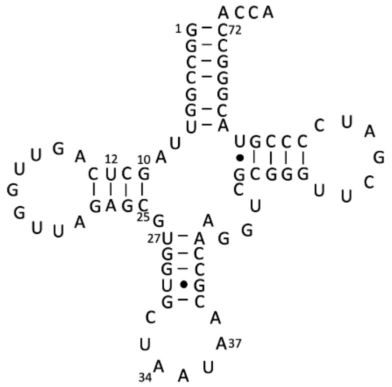


**Figure S1. hKEOPS - tRNA$^{Ile}_{AAU}$ interaction followed by fluorescence anisotropy**

(A) 3'-fluorescein-labeled tRNA$^{Ile}_{AAU}$ (40 nM) was titrated with increasing concentration of hKEOPS in buffer containing 4 mM, 50 mM or 200 mM of NaCl. Saturation curves have been fitted to a rectangular hyperbolic model using Prism10 software.

(B) Table summarizing the data fitting with equilibrium dissociation constant $K_D$ in nM.

(C) Clover-leaf representation of human tRNA$^{Ile}_{AAU}$

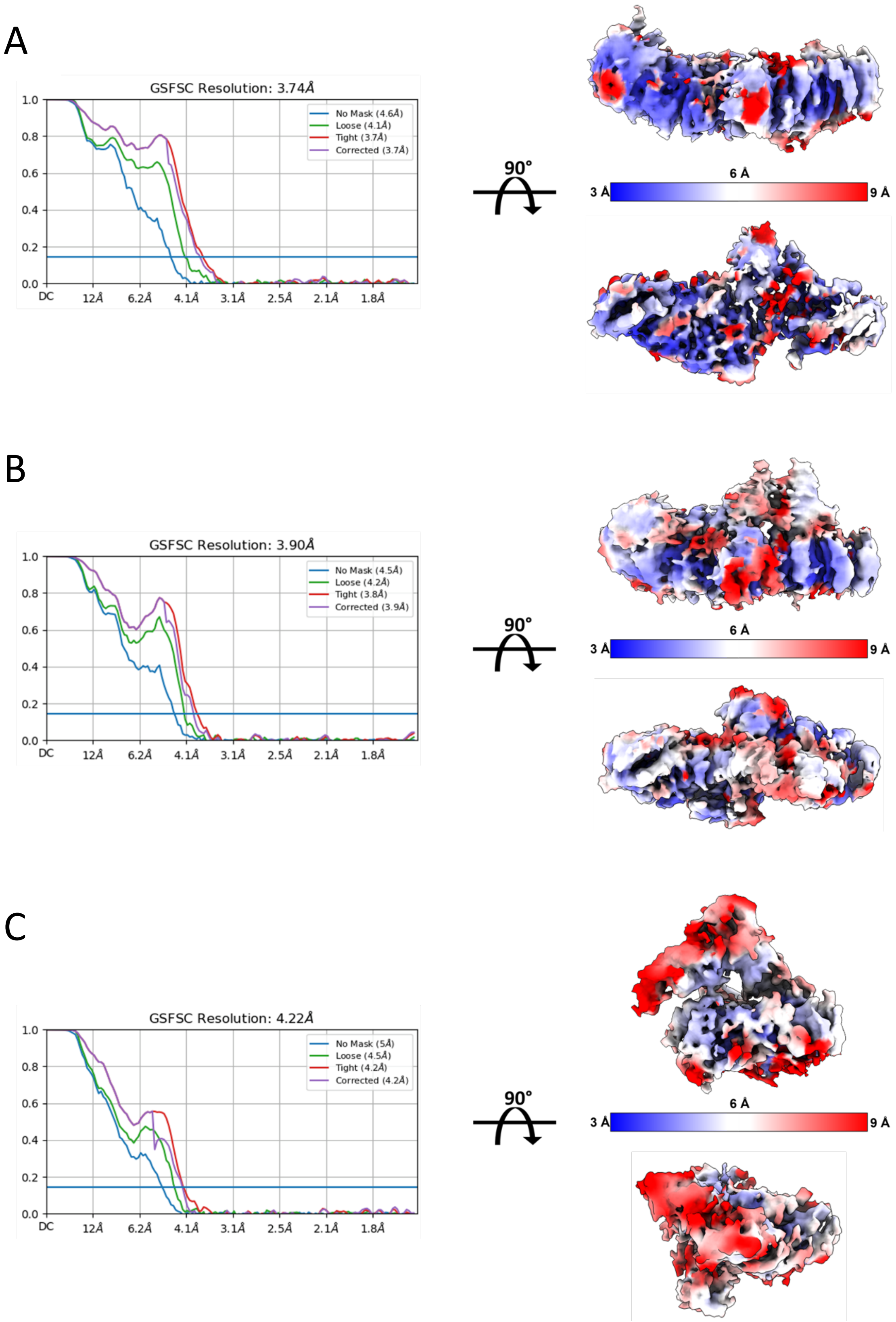
A
GSFSC Resolution: 3.74Å
No Mask (4.6Å)
Loose (4.1Å)
Tight (3.7Å)
Corrected (3.7Å)
1.0
0.8
0.6
0.4
0.2
0.0
DC 12Å 6.2Å 4.1Å 3.1Å 2.5Å 2.1Å 1.8Å
90°
3 Å
6 Å
9 Å
B
GSFSC Resolution: 3.90Å
No Mask (4.5Å)
Loose (4.2Å)
Tight (3.8Å)
Corrected (3.9Å)
1.0
0.8
0.6
0.4
0.2
0.0
DC 12Å 6.2Å 4.1Å 3.1Å 2.5Å 2.1Å 1.8Å
90°
3 Å
6 Å
9 Å
C
GSFSC Resolution: 4.22Å
No Mask (5Å)
Loose (4.5Å)
Tight (4.2Å)
Corrected (4.2Å)
1.0
0.8
0.6
0.4
0.2
0.0
DC 12Å 6.2Å 4.1Å 3.1Å 2.5Å 2.1Å 1.8Å
90°
3 Å
6 Å
9 Å

D

| | apo-hKEOPS | hKEOPS/tRNA | hKEOPS subcomplex/tRNA |
|---|---|---|---|
| **Data collection and processing** | | | |
| Microscope | Titan Krios | | |
| Camera | Falcon 4i | | |
| Magnification | 165,000 | | |
| Voltage | 300 kV | | |
| Defocus range (µm) | -1 to -3 (0.2 steps) | | |
| Pixel size | 0.77 Å/pixel | | |
| Total exposure | 40 $e^-/Å^2$ | | |
| Movies collected | 14,328 | | |
| Movies used | 13,411 | | |
| Initial number of particles | 9,613,829 | | |
| Final number of particles | 214,527 | 214,607 | 121,547 |
| Final map resolution | 3.74 Å | 3.91 Å | 4.22 Å |
| **Refinement** | | | |
| Model composition | | | |
| Non-hydrogen atoms | 6,779 | 8,422 | 6,086 |
| Protein residues | 873 | 873 | 567 |
| RNA residues | 0 | 77 | 77 |
| B-factors (Å2) min/max/mean | | | |
| Protein | 47.27/212.14/124.29 | 12.36/116.82/68.17 | 0.98/110.04/44.71 |
| Nucleotide | - | 86.80/198.77/135.40 | 27.15/189.40/99.07 |
| R.m.s deviation | | | |
| Bond lengths (Å) | 0.004 | 0.004 | 0.003 |
| Bond angles (°) | 0.820 | 0.867 | 0.704 |
| Model vs. Data | | | |
| Cross Correlation (mask) | 0.61 | 0.49 | 0.56 |
| Cross Correlation (box) | 0.70 | 0.65 | 0.67 |
| Cross Correlation (peaks) | 0.50 | 0.39 | 0.43 |
| Cross Correlation (volume) | 0.60 | 0.46 | 0.53 |
| **Validation** | | | |
| MolProbity score | 2.37 | 2.46 | 2.15 |
| Clash score | 20.25 | 27.98 | 17.43 |
| Ramachandran plot (%) | | | |
| Outliers | 0.00 | 0.00 | 0.00 |
| Allowed | 10.54 | 9.04 | 6.06 |
| Favored | 89.46 | 90.96 | 93.94 |
| Rotamer outliers (%) | 0.14 | 0.14 | 0.00 |
| Cβ outliers (%) | 0.00 | 0.00 | 0.00 |

**Figure S2. Global and local resolution of the cryo-EM maps and statistics.**
Left panels: Gold-standard Fourier shell correlation (GSFSC) curves of the final maps, as calculated by CryoSPARC. Right panels: local resolution maps calculated using CryoSPARC. The color gradient indicates local resolution, ranging from 3 Å (blue) to 9 Å (red).
(A) Resolution estimation of the apo-hKEOPS cryo-EM map.
(B) Resolution estimation of the hKEOPS/tRNA complex cryo-EM map.
(C) Resolution estimation of the hKEOPS/tRNA sub-complex.
(D) Cryo-EM data collection, refinement and validation statistics.

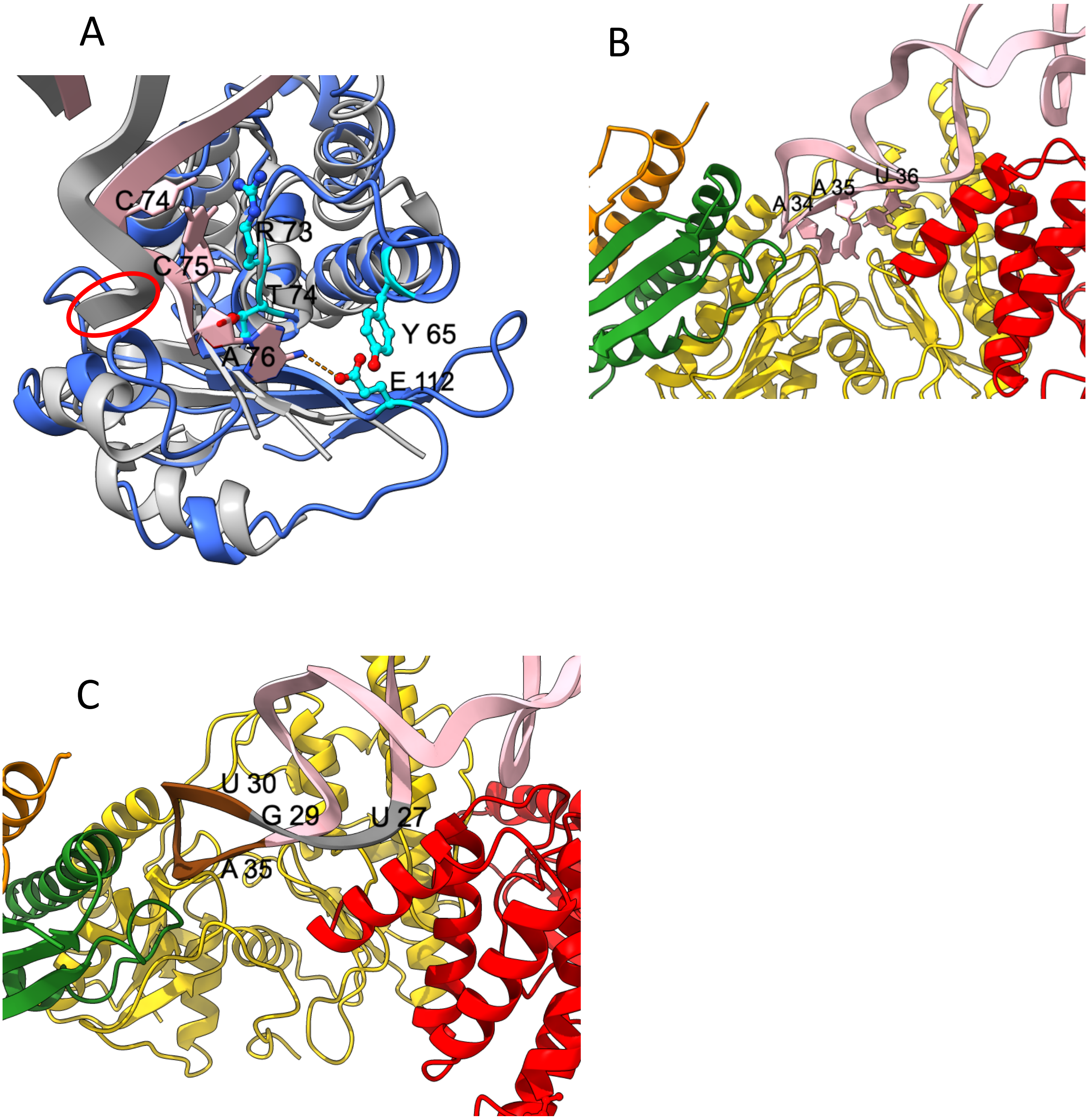


**Figure S3. Interaction of hKEOPS with tRNA$^{Ile}_{AAU}$.**

(A) Superposition of the CCA tails in archaeal Cgi121 and human TPRKB tRNA-bound complexes. The Cgi121/tRNA complex is shown in grey, TPRKB in blue, and tRNA$^{Ile}_{AAU}$ in pink. The CCA tail of tRNA is highlighted with a red circle. Y65$^{TPRKB}$, R73$^{TPRKB}$, T74$^{TPRKB}$ and E112$^{TPRKB}$ are in cyan ball-and-stick.

(B) View of the AC-loop region of tRNA in the hKEOPS– tRNA$^{Ile}_{AAU}$ complex, showing bases $A_{34}$–$A_{35}$–$U_{36}$.

(C) The AC-loop region of tRNA is interacting with LAGE3 via the loop $U_{30}$–$A_{35}$ (brown), and with TP53RK C-ter $\alpha$-helix via its $U_{27}$–$G_{29}$ segment (grey).

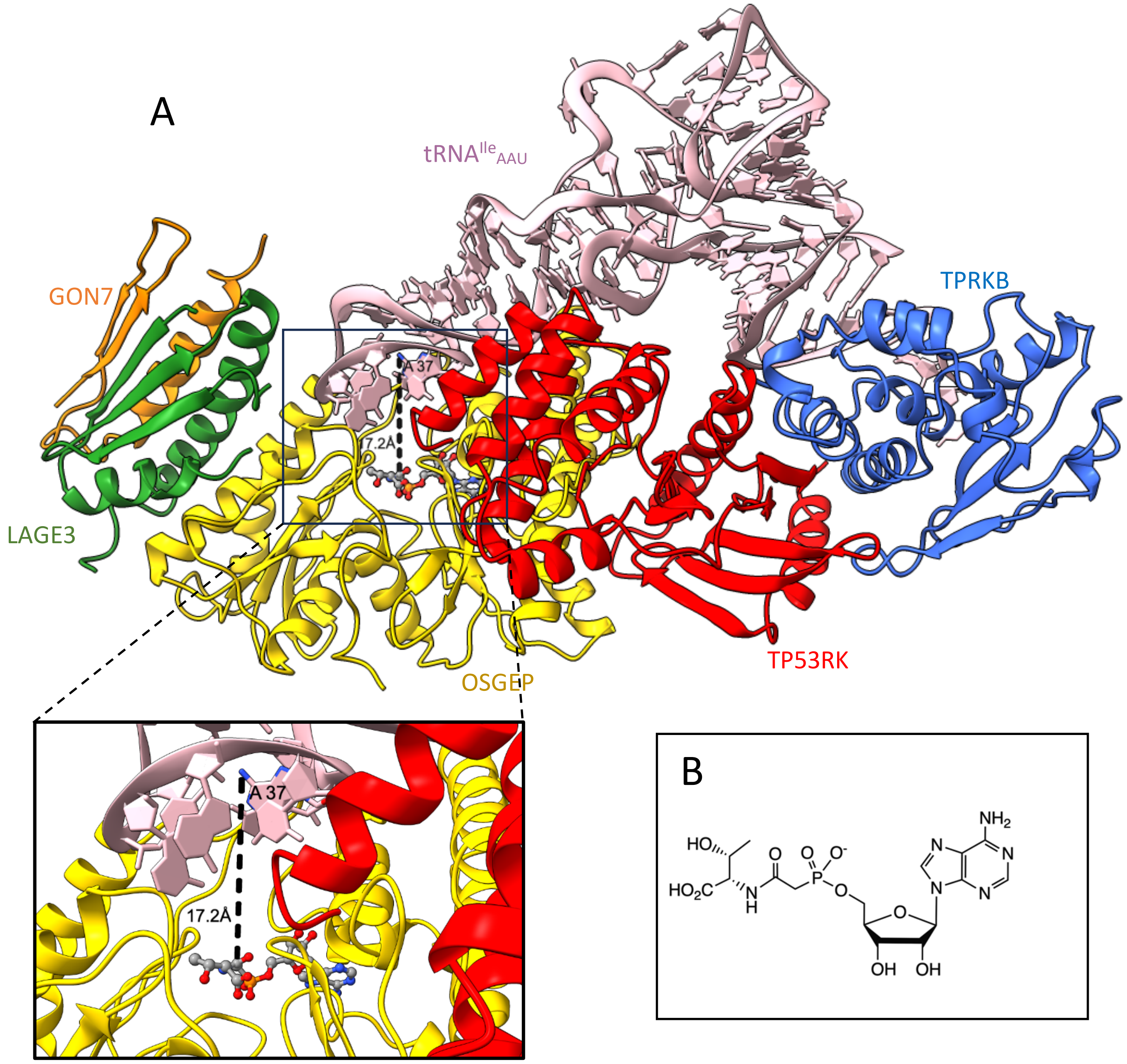


**Figure S4. Superposition of hKEOPS/tRNA with the bacterial TsaD subunit bound to a TC-AMP analog.**
(A) hKEOPS/tRNA was superimposed onto the bacterial TsaD bound to BK951 (PDB ID: 6Z81), where BK951 mimics the second substrate (TC-AMP) of the $t^6A$ modification reaction. For clarity, the TsaD structure is omitted; only the BK951 molecule is shown (ball-and-stick representation).
The inset below shows a close-up of the OSGEP active site, highlighting that the $N^6$ atom of the tRNA substrate base $A_{37}$ is positioned 17 Å from the carbamoyl group of the TC-AMP analog. This distance indicates that the tRNA anticodon loop is not correctly positioned to support catalysis.
(B) BK951 (TC-AMP anolog) structure.

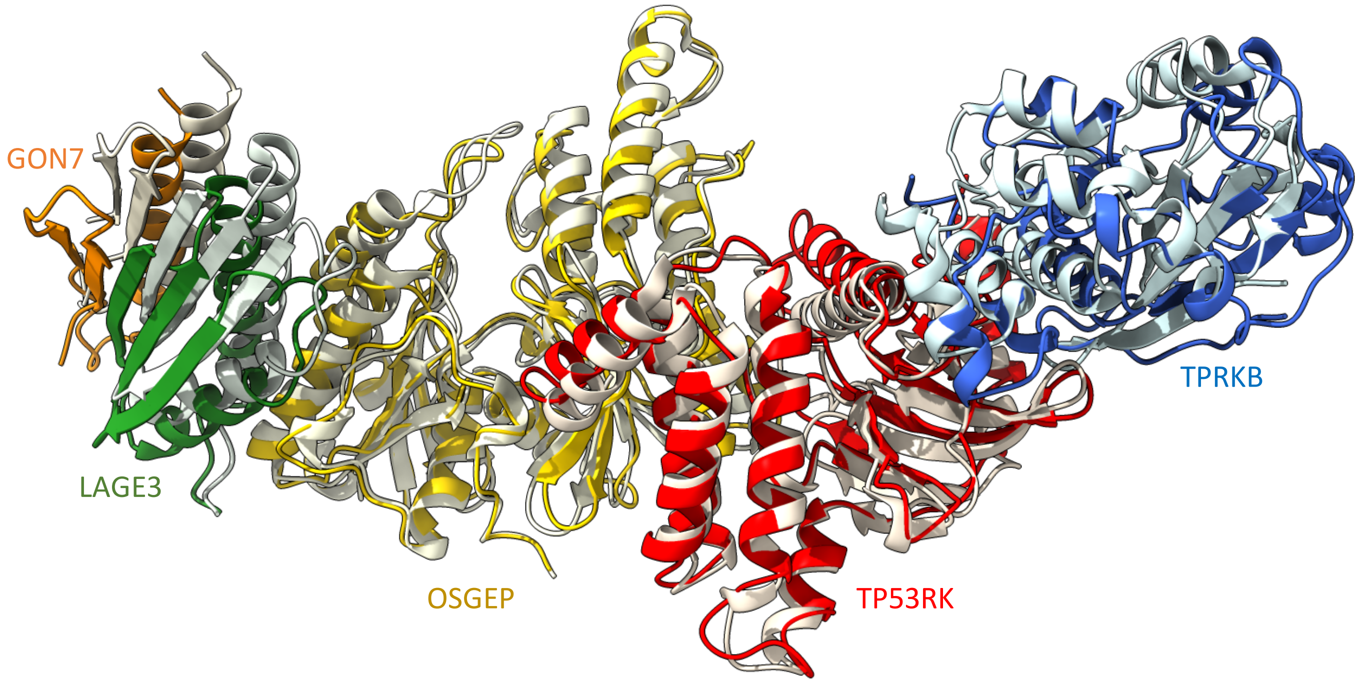


**Figure S5. Superposition of apo-hKEOPS with crystal structures of hKEOPS subcomplexes.**
The apo-hKEOPS structure is superimposed with the crystal structures of the OSGEP/LAGE3/GON7 subcomplex (PDB ID: 6GWJ) and the TP53RK/TPRKB subcomplex (PDB ID: 6WQX). Subunit colors are bright for apo-hKEOPS and lighter for the corresponding subcomplexes: GON7 (orange), LAGE3 (green), OSGEP (yellow), TP53RK (red), and TPRKB (blue).

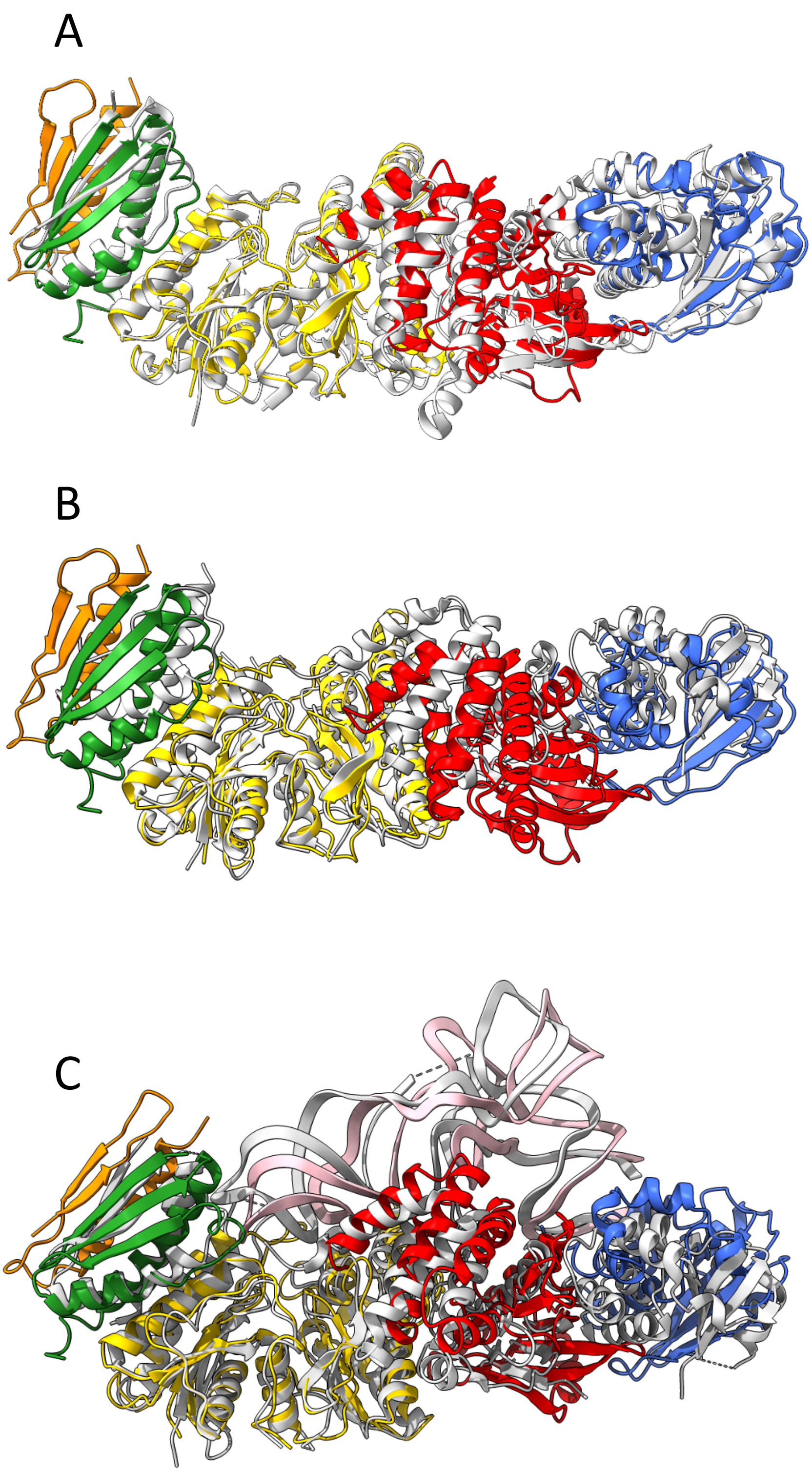

**Figure S6. Superposition of KEOPS complexes.**
Subunits of hKEOPS are colored as follows: GON7 (orange), LAGE3 (green), OSGEP (yellow), TP53RK (red), and TPRKB (blue).
(A) Superposition of apo-hKEOPS with *Arabidopsis thaliana* apo-KEOPS (AtKEOPS; PDB ID: 8K20, shown in grey).
(B) Superposition of apo-hKEOPS with archaeal apo-KEOPS (PDB ID: 8UNK, shown in grey).
(C) Superposition of the hKEOPS/tRNA complex with the archaeal KEOPS/tRNA complex (PDB ID: 8UP5, shown in grey).

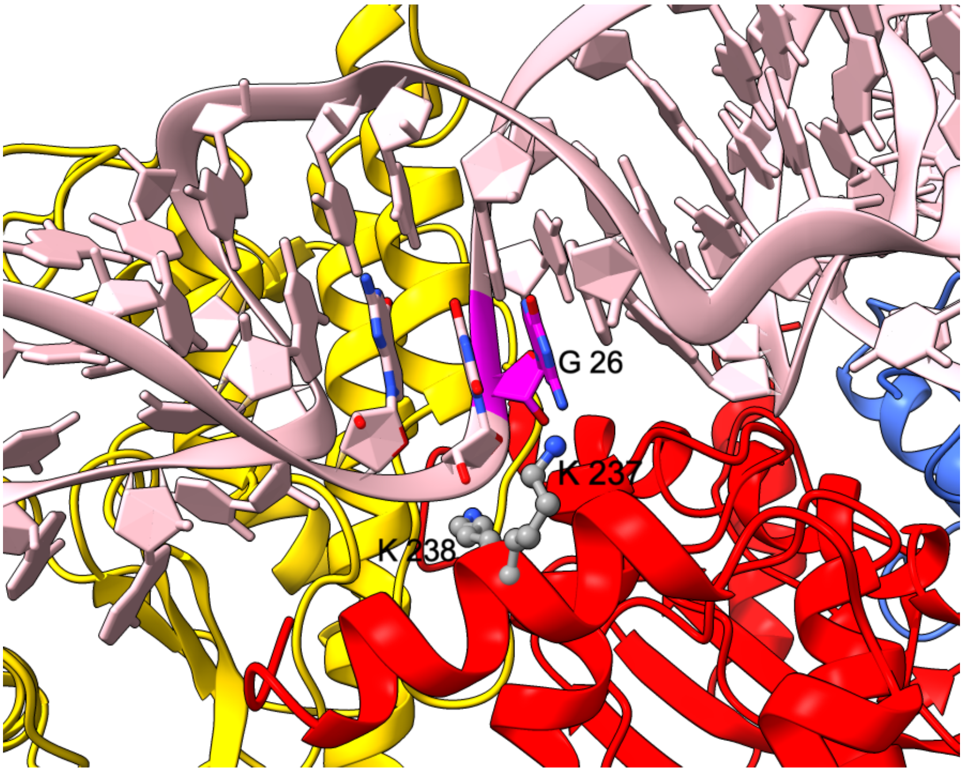


**Figure S7. Interactions between the C-terminal α-helix of TP53RK and the tRNA$^{Ile}_{AUU}$ backbone.**
The C-terminal α-helix of TP53RK interacts with the sugar-phosphate backbone of nucleotides $G_{26}U_{27}G_{28}$ of tRNA$^{Ile}_{AUU}$. Notably, base $G_{26}$ (shown in purple) remains stacked within the core of the tRNA structure.

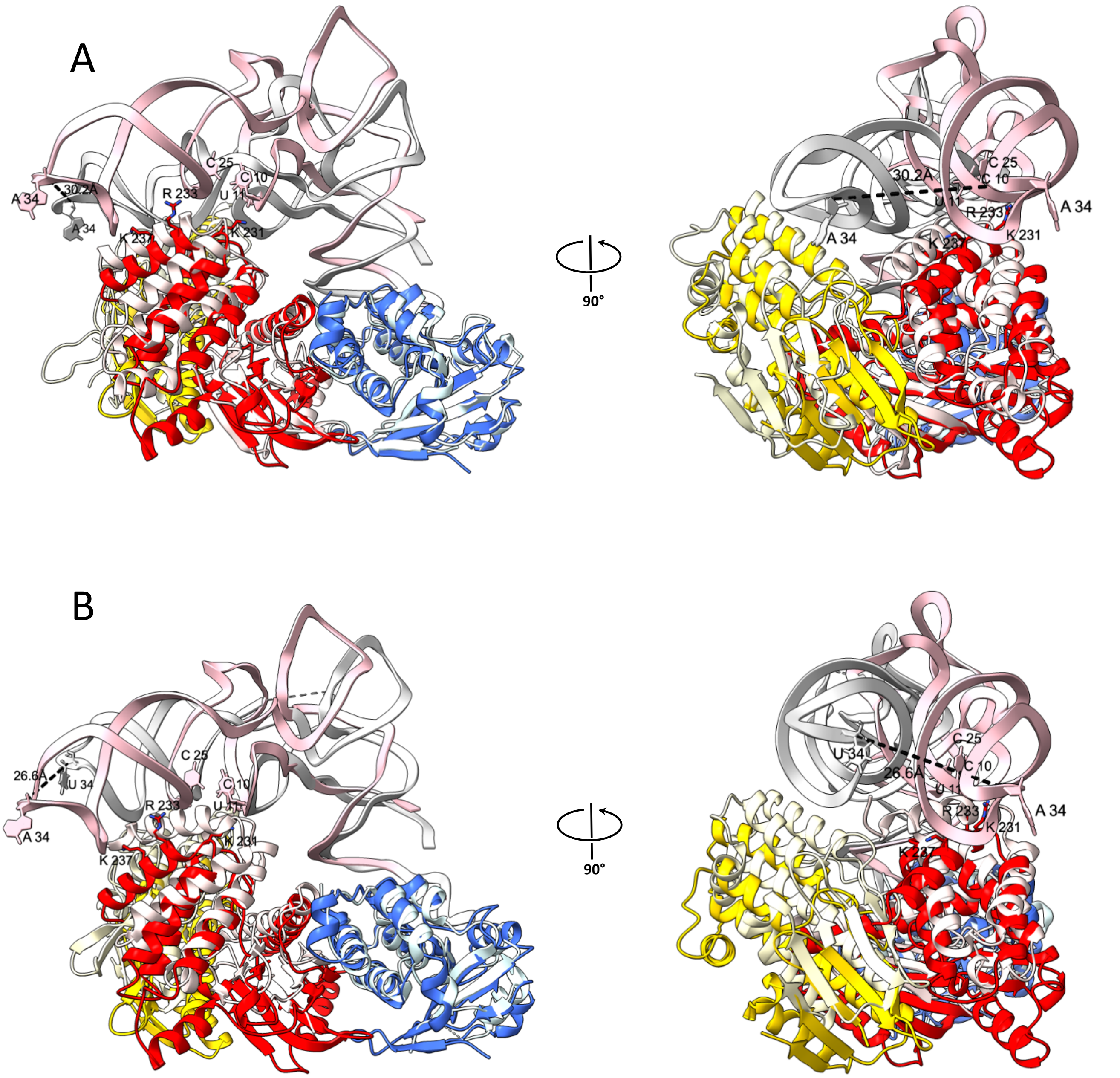


**Figure S8. Comparison of the hKEOPS subcomplex and full KEOPS complexes.**

(A) Superposition of the hKEOPS/tRNA subcomplex (tRNA in pink) with the full hKEOPS/tRNA complex (tRNA in grey). Subunits are colored brightly for the subcomplex and lightly for the full complex: OSGEP (yellow), TP53RK (red), and TPRKB (blue). A displacement of more than 30 Å is observed for the anticodon (AC) loop of tRNA toward the TP53RK subunit in the human subcomplex. For clarity, GON7, LAGE3, and portions of OSGEP (residues 1–125 and 288–335) are omitted from the full hKEOPS complex.

(B) Superposition of the hKEOPS/tRNA subcomplex (tRNA in pink) with the archaeal KEOPS/tRNA complex (tRNA in grey; PDB-ID 8UP5). Subunit colors follow the same scheme as in panel A: bright for the human subcomplex and light for the archaeal complex. A similar >26 Å shift of the AC-loop toward TP53RK is observed. For clarity, Pcc1, LAGE3, and portions of Kae1 (residues 1–125 and 276–324) are omitted.

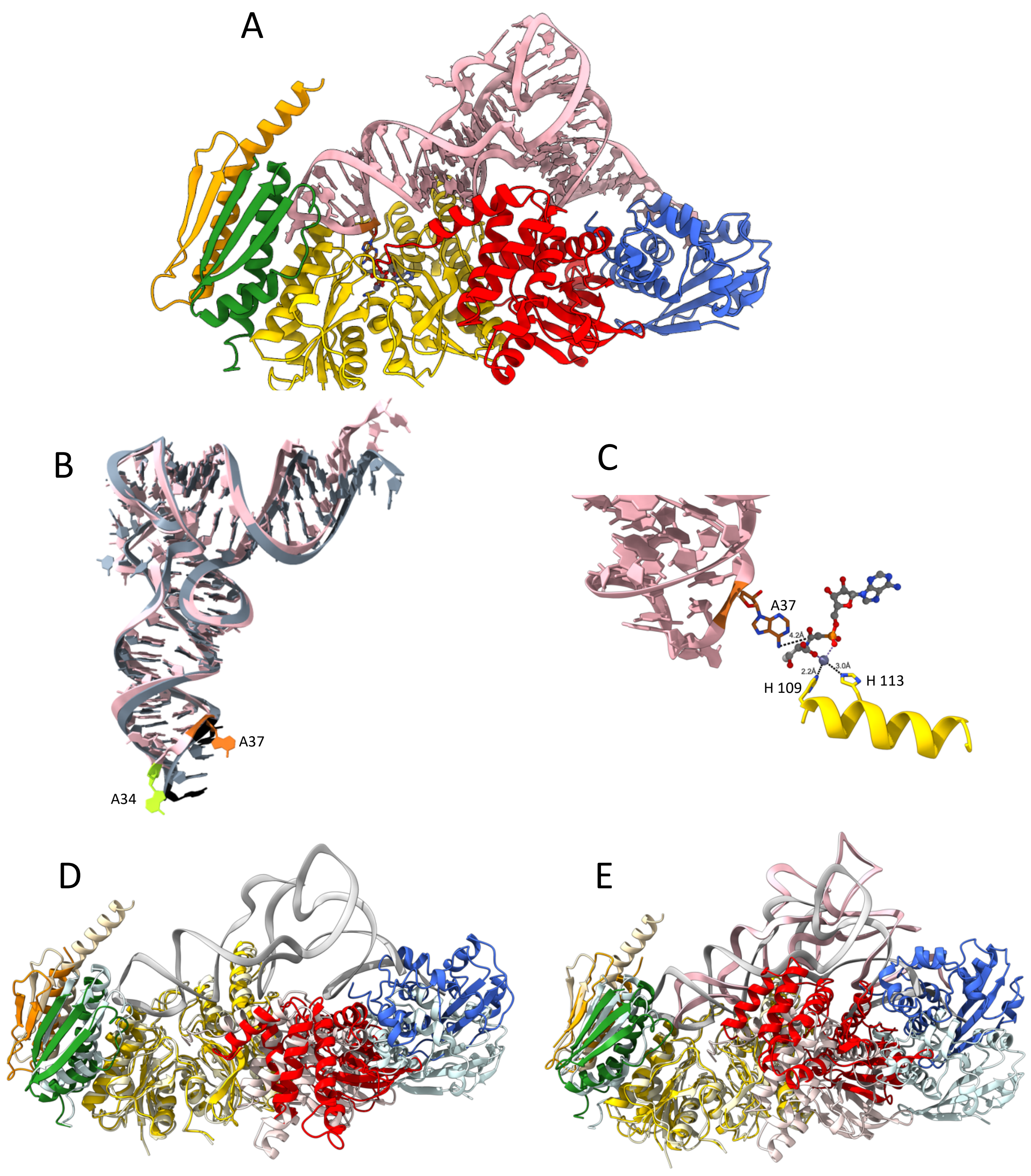


**Figure S9. AF3 models of hKEOPS/tRNA complexes.**
Color code: orange : GON7; green LAGE3 ; yellow : OSPEG; red: TP53RK; blue: TPRKB; pink: tRNA
For clarity, disordered predicted segments (plDDT below 70) of LAGE3 (1-58), GON7 (61-100) and TP53RK (1-15) in the AF3 model are omitted.
(A) AF3 model of hKEOPS bound to tRNA$^{Ile}$AUU and superposed to the TC-AMP analog (i.e. BK951 molecule, ball-and-stick, PDB-ID: 6Z81) within OSGEP active site.
(B) Superposition of predicted AF3 model of cognate tRNA$^{Ile}_{AUU}$ alone (grey, showing base 34 and 37 in black that are folded in the AC-loop) or in the presence of hKEOPS (pink, showing base 34 in green and base 37 in orange that are flipped outside the AC-loop).

(C) A zoom view of the active site of (A) with a slightly different orientation in order to emphasize the proximity of base A37 (brown) to the carbonyl of TC-AMP analog. Conserved $H109^{OSGEP}$ and $H113^{OSGEP}$ close to the $Zn^{2+}$ metal ion (grey) are shown in yellow sticks.
(D) Superposition of apo-hKEOPS to AF3 model of hKEOPS/tRNA (light color and grey tRNA).
(E) Superposition of tRNA-bound cryo-EM structure of hKEOPS to AF3 tRNA-bound model of hKEOPS (light color and grey tRNA).

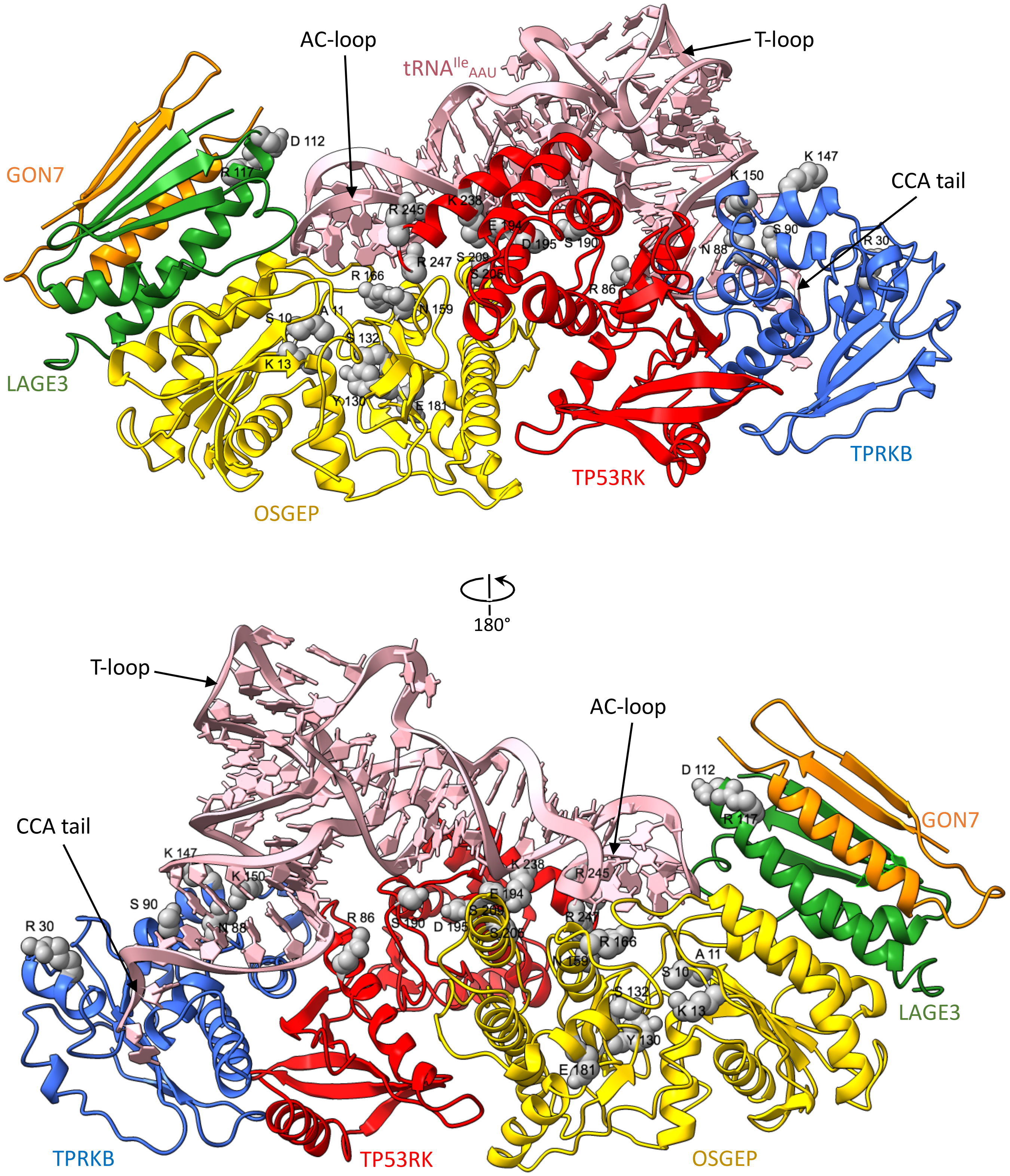


**Figure S10. Positions of residues mutated to validate the cryo-EM structure.**

Residues targeted for mutagenesis are shown as grey spheres mapped onto the cryo-EM structure of hKEOPS/tRNA$^{Ile}_{AAU}$ .

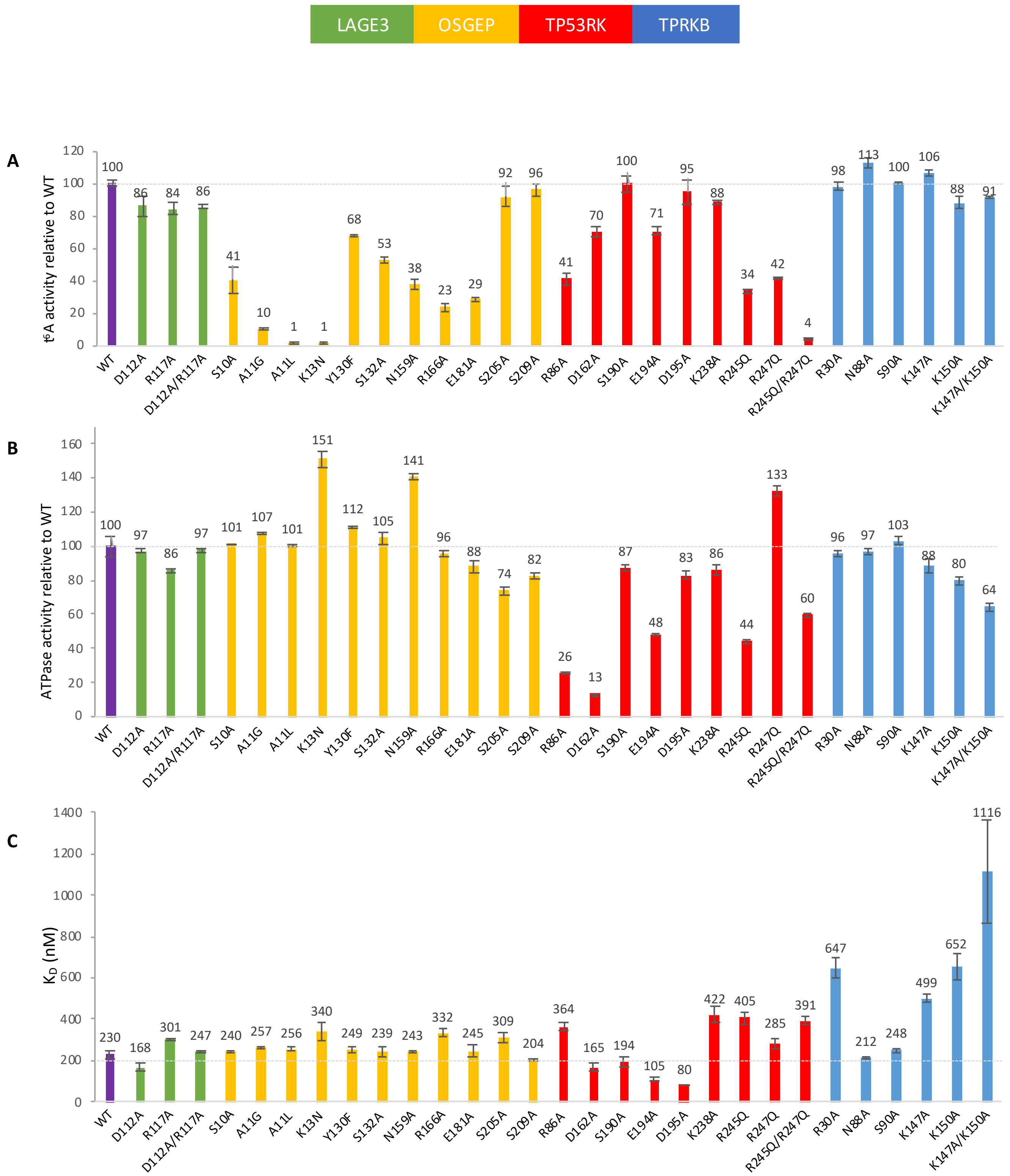


**Figure S11. Biochemical characterization of mutants designed to probe the cryo-EM structure of hKEOPS/tRNA$^{Ile}_{AAU}$.**

Mean values and standard deviations are shown for each mutant.

(A) t$^6$A modification activity, expressed relative to WT hKEOPS.

(B) ATPase activity expressed relative to WT hKEOPS (see supplementary methods).

(C) Dissociation constants ($K_D$) for the interaction between hKEOPS mutants and tRNA$^{Ile}_{AAU}$ (see supplementary methods).

The D162A$^{TP53RK}$ is a catalytic mutant of the ATPase activity

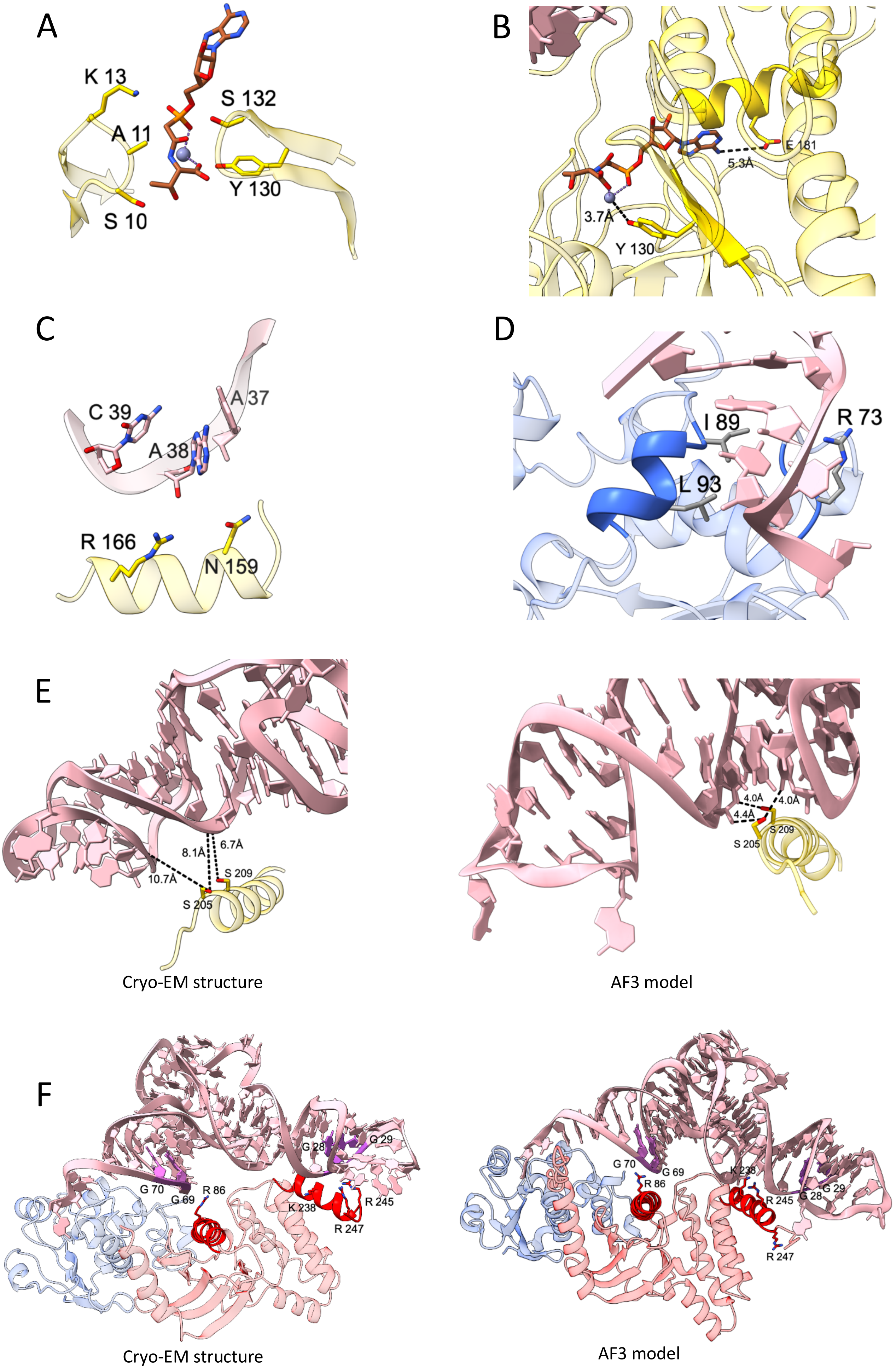
A
K 13
A 11
S 132
Y 130
S 10
B
E 181
5.3Å
3.7Å
Y 130
C
C 39
A 37
A 38
R 166
N 159
D
I 89
R 73
L 93
E
8.1Å
6.7Å
10.7Å
S 209
S 205
Cryo-EM structure
4.0Å
4.0Å
4.4Å
S 209
S 205
AF3 model
F
G 28
G 29
G 70
G 69
R 86
K 238
R 245
R 247
Cryo-EM structure
G 70
G 69
R 86
K 238
R 245
G 28
G 29
R 247
AF3 model

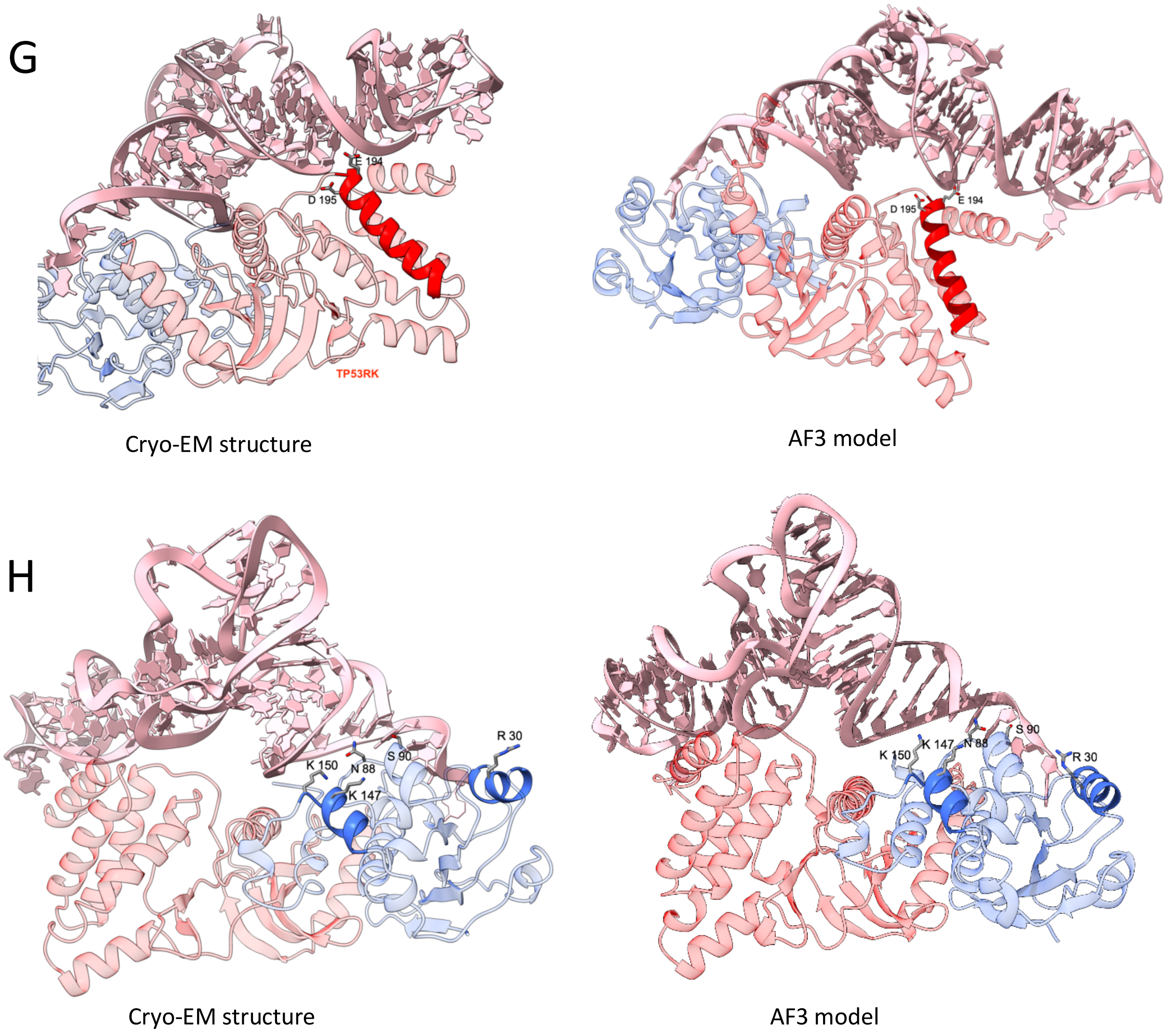


**Figure S12. Mutations sampling of the OSGEP active site and the hKEOPS tRNA interface.**

(A) Close-up of the OSGEP active site with BK951 (TC-AMP analog; brown) modelized from PDB-ID: 6Z81 showing the tested mutations S10A$^{OSGEP}$, A11L$^{OSGEP}$, K13N$^{OSGEP}$, S132A$^{OSGEP}$ and Y130F$^{OSGEP}$.

(B) Close-up on E181$^{OSGEP}$ and Y130$^{OSGEP}$ showing the proximity with TC-AMP analog (brown) modelized from PDB-ID: 6Z81.

(C) Close-up on N159$^{OSGEP}$ and R166$^{OSGEP}$ (yellow) and bases 37-39 of tRNA (pink).

(D) Mutations at R73$^{TPRKB}$, I89$^{TPRKB}$ and L93$^{TPRKB}$ that have been shown to lose affinity for tRNA are in contact to CCA-tail in the Cryo-EM structure

(E) Left**:** close-up on S205$^{OSGEP}$ and S209$^{OSGEP}$ (yellow) and tRNA (pink) of the hKEOPS/tRNA structure. Right: same as left view for the AF3 model, showing that the $\alpha$-helix has shifted toward the D-arm of tRNA in the AF3 model.

(F) Left: interaction of TP53RK (red) with tRNA (pink and light purple bases) in the hKEOPS/tRNA structure, showing positively charged residues on $\alpha$-helices (dark red) near the tRNA backbone. Right: the same view in the AF3 model, where R247 is exposed to solvent and does not interact with tRNA.

(G) Left: interaction of TP53RK (red) with tRNA (pink) showing the negatively charged E194$^{TP53RK}$ and D195$^{TP53RK}$ on $\alpha$-helix (dark-red) nearby the tRNA backbone in the hKEOPS/tRNA structure. right: the same view in the AF3 model.

(H) Left: interaction of TPRKB (blue) with tRNA (pink) highlighting the $\alpha$-helices (dark blue) containing positively charged R30$^{TPRKB}$ near the CCA-tail of tRNA, as well as K147$^{TPRKB}$ and K150$^{TPRKB}$ near the acceptor stem. right: same as left with AF3 predicted model showing R30 in a different orientation and closer to CCA-tail.

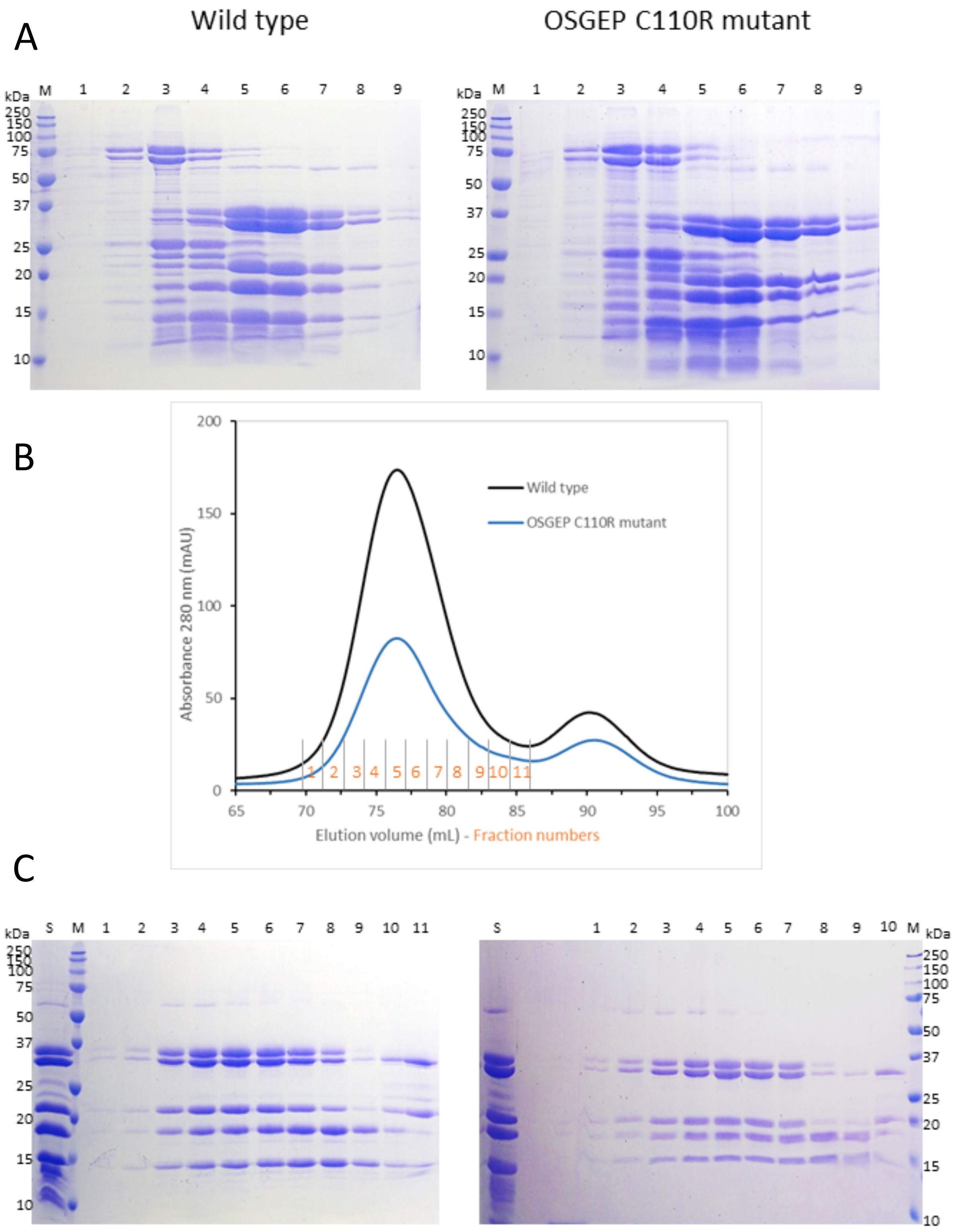


**Figure S13. Purification of hKEOPS**.

Expression and purification of recombinant proteins protocol can be found in the supplementary methods section. A) SDS-PAGE analysis of fractions from the Ni-NTA affinity chromatography step. Lane M: protein molecular weight markers. Lanes 1–9: fractions eluted with 60, 80, 100, 200 (two lanes), and 400 mM (four lanes) imidazole, respectively. WT hKEOPS is shown on the left; hKEOPS carying the C110R$^{OSGEP}$ mutant on the right.

(B) Gel filtration chromatography profiles of WT hKEOPS and hKEOPS carying the C110R$^{OSGEP}$ mutant .

(C) SDS-PAGE analysis of gel filtration fractions. Lane S: loaded sample (pooled from Ni-NTA fractions corresponding to lanes 5–8). Lanes 2–6: eluted fractions containing hKEOPS.

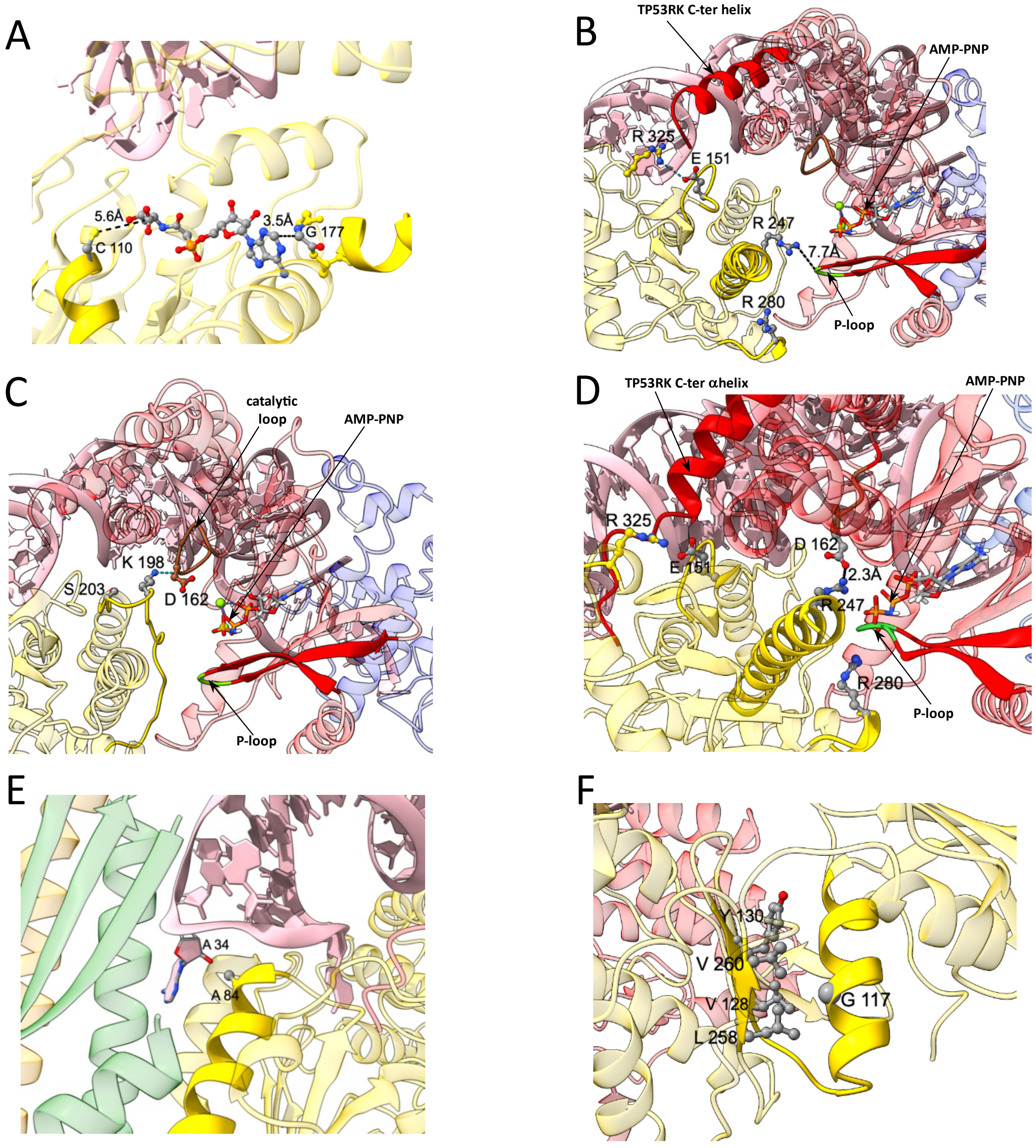


**Figure S14. GAMOS mutations in the OSGEP subunit.**

(A) Model of a TC-AMP analog bound in the active site of OSGEP, showing the proximity of residues C110 and G177.

(B) View of the interface between OSGEP (yellow) and TP53RK (red). An AMP-PNP molecule (shown in ball-and-stick representation) is modeled in the TP53RK active site using PDB ID: 6WQX (Li J. et al., 2021). A salt bridge between E151$^{\text{OSGEP}}$ and R325$^{\text{OSGEP}}$ is indicated by a blue dotted line.

(C) Hydrogen bond between K198$^{\text{OSGEP}}$ and G161$^{\text{TP53RK}}$ (dotted line), located near the catalytic residue D162$^{\text{TP53RK}}$

(D) OSGEP–TP53RK interface from the AF3-predicted hKEOPS/tRNA complex, showing the proximity of R247$^{\text{OSGEP}}$ to both D162$^{\text{TP53RK}}$ and the $\gamma$-phosphate of AMP-PNP.

(E) Position of residue A84$^{\text{OSGEP}}$ (a GAMOS mutant site) in the AF3 model, near the ribose of tRNA base A$_{34}$.

(F) Mutation site G117$^{\text{OSGEP}}$ facing the hydrophobic environment of the adjacent $\beta$-sheet

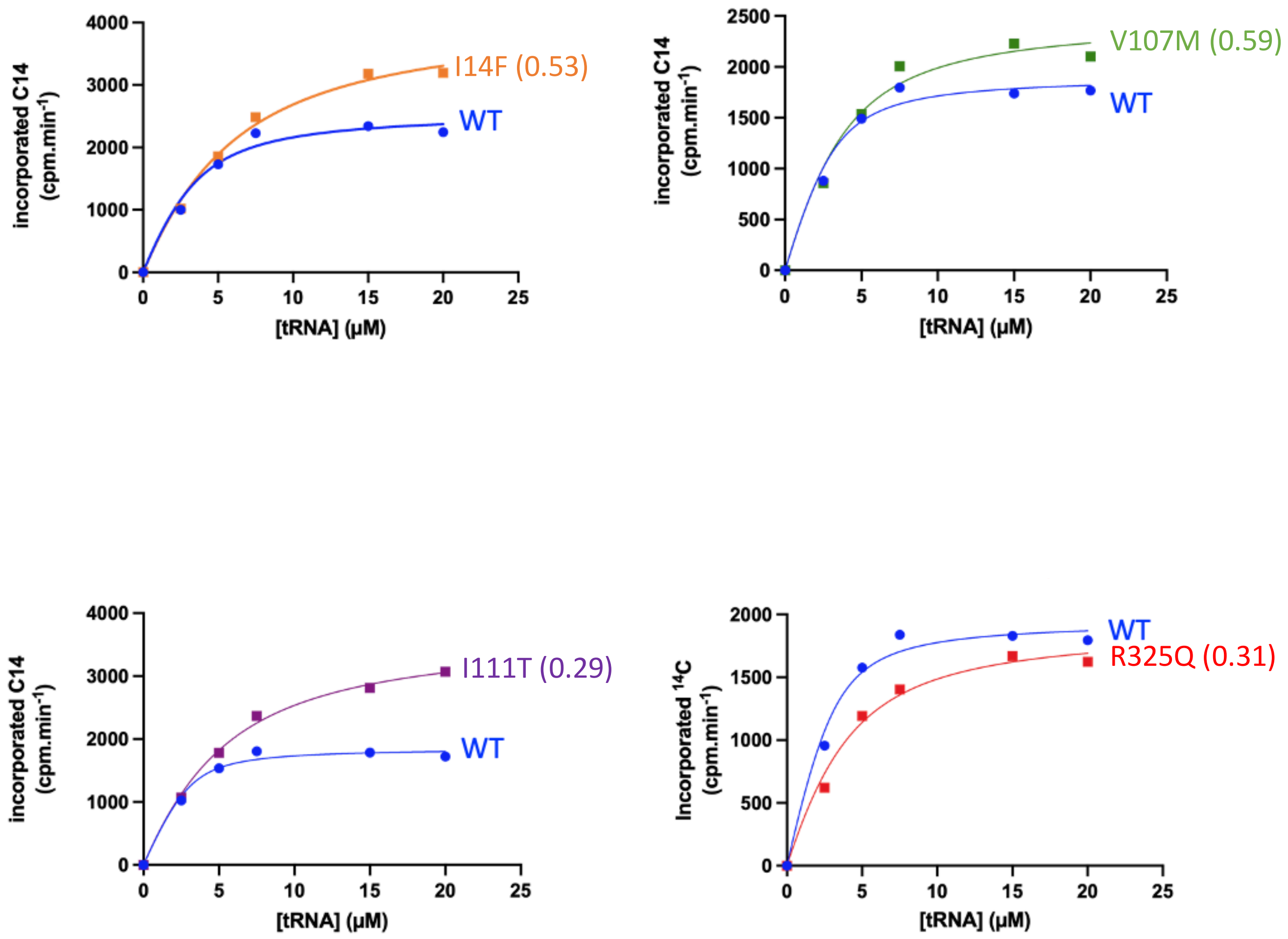


**Figure S15. Michaelis-Menten saturation curves of GAMOS mutants and WT hKEOPS.**
I14F$^{OSGEP}$, I11T$^{OSGEP}$, and V107M$^{OSGEP}$ are mutants with higher-than-WT activity while R325Q$^{OSGEP}$ has a reduced activity.
Catalytic efficiency (kcat/$K_M$) ratios of each mutant relative to WT are shown in parentheses.

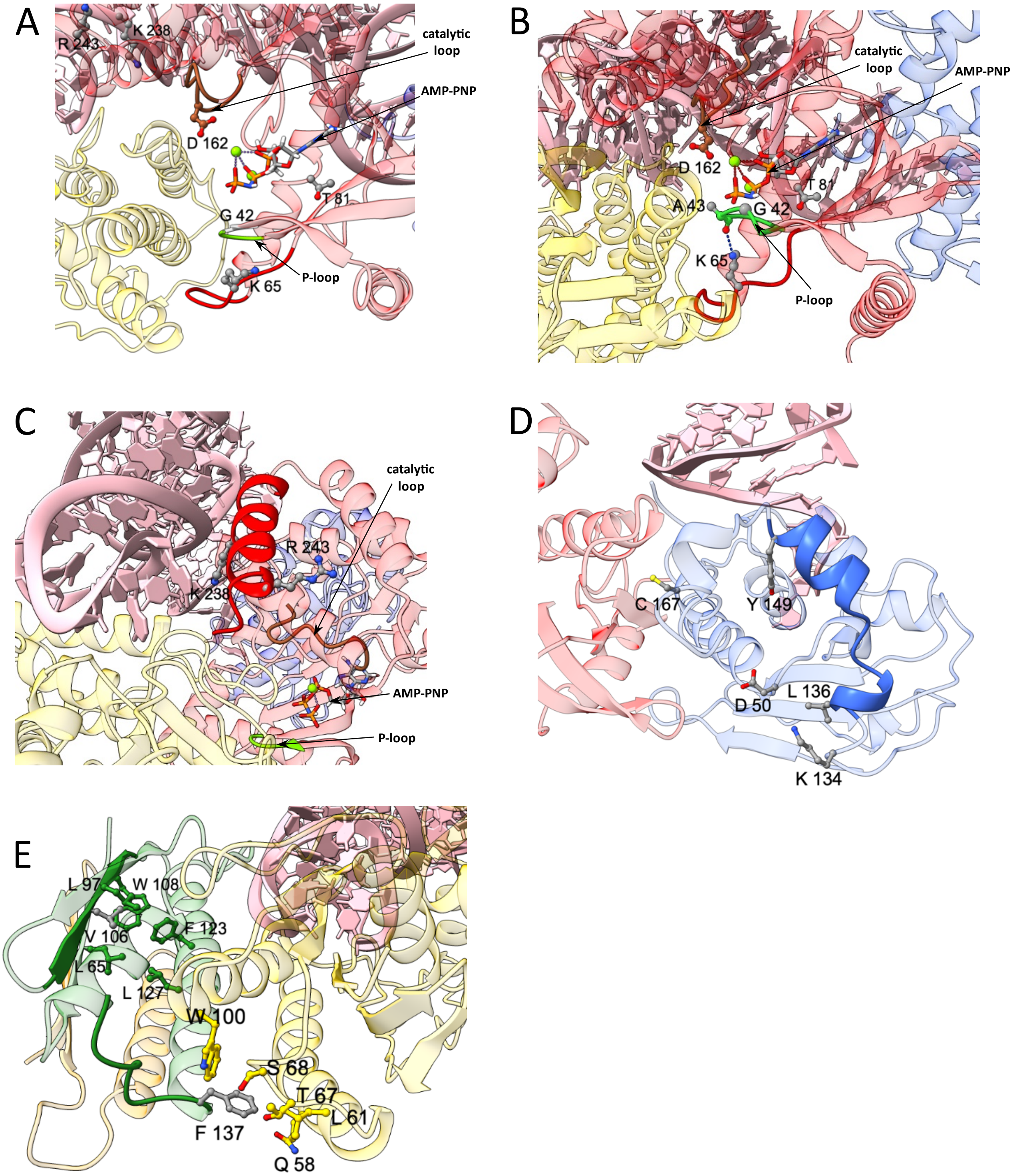


**Figure S16. GAMOS mutations in the TP53RK, TPRKB, and LAGE3 subunits.**

(A) View of the TP53RK active site, highlighting GAMOS-associated mutations: $G42^{TP53RK}$, $K65^{TP53RK}$ and $T81^{TP53RK}$. The catalytic loop is shown in brown and the P-loop in green. The catalytic residue $D162^{TP53RK}$ and AMP-PNP (shown as sticks) are also indicated.

(B) View of the TP53RK active site from the AF3 model of the hKEOPS/tRNA complex. The side chain of $K65^{TP53RK}$ forms a hydrogen bond with the carbonyl group of $A43^{TP53RK}$ (blue dotted line).

(C) View of the TP53RK C-terminal helix and its interaction with tRNA.

(D) Locations of selected GAMOS mutations within the TPRKB subunit.

(E) Locations of selected GAMOS mutations within the LAGE3 subunit, including $L106^{LAGE3}$ and $F137^{LAGE3}$ (shown in grey), alongside surrounding side chains from LAGE3 (green) and OSGEP (yellow).

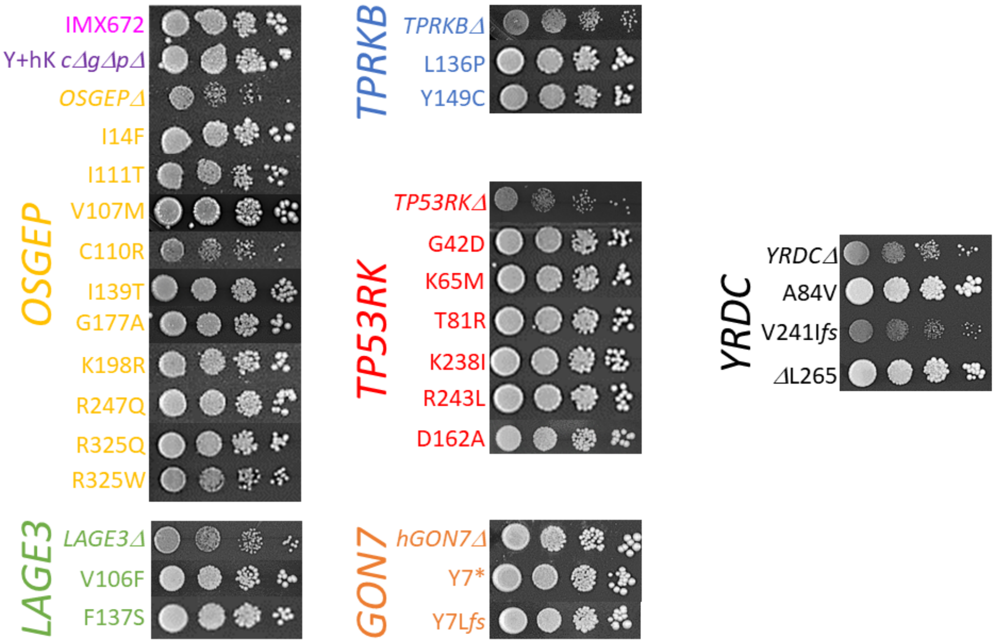


**Figure S17. Effect of GAMOS-associated mutations on the fitness of the fully humanized yeast strain.**
The effects of individual GAMOS-associated point mutations in each human gene of the $t^6A$ biosynthesis pathway were evaluated using a fitness assay on plate. The color code is used to identify the hKEOPS subunits and their corresponding mutations. Growth profiles of the intact yeast recipient strain IMX672 (ySua5 + yKEOPS) and the fully humanized yeast strain Y+hK *cΔgΔpΔ* (YRDC + hKEOPS *cgi121Δ gon7Δ pcc1Δ*) are shown as normal growth controls. For reference, the phenotypes resulting from individual deletions of the human $t^6A$ pathway genes are also displayed, allowing direct comparison with their respective GAMOS-associated mutant alleles. $D162A^{TP53RK}$, a non-GAMOS mutation that abolishes the ATPase activity of TP53RK, is included as a control for functional loss.

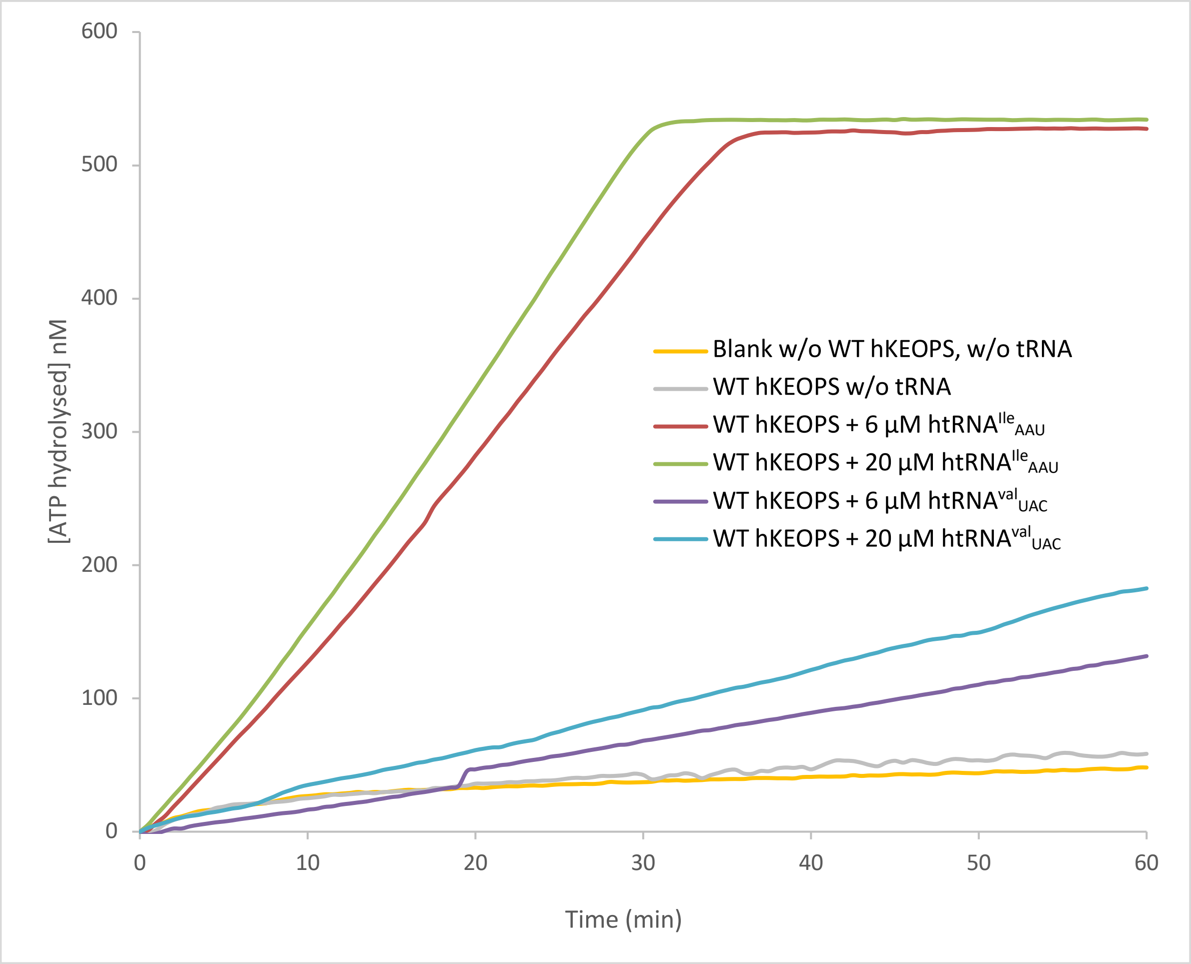


**Figure S18. Time course of ATP hydrolysis by WT hKEOPS using cognate and non-cognate tRNA substrates.** ATPase activity (see supplementary methods) was measured over time for WT hKEOPS in the absence of tRNA (grey line), in the presence of cognate tRNA$^{Ile}_{AAU}$ transcript at 6 µM (red line) or 20 µM (green line), and in the presence of non-cognate tRNA$^{Val}_{UAC}$ transcript at 6 µM (purple line) or 20 µM (blue line).

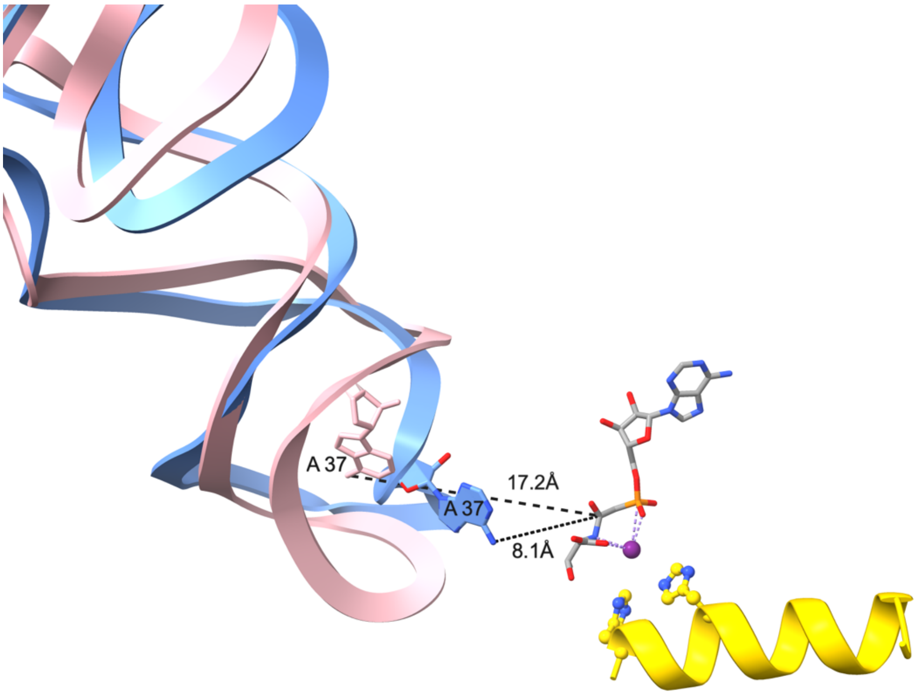


**Figure S19. Distinct conformations of the tRNA anticodon loop in hKEOPS and ADAT2/3 desaminase.** the tRNA of the cryo-EM structure of the ADAT2/3–tRNA complex (blue, PDB ID : 8AW3) has been superimposed to our hKEOPS model of Fig S4 (pink). The anticodon loop of ADAT2/3 adopts a more open conformation, exposing base 37 closer to the TCAMP intermediate.

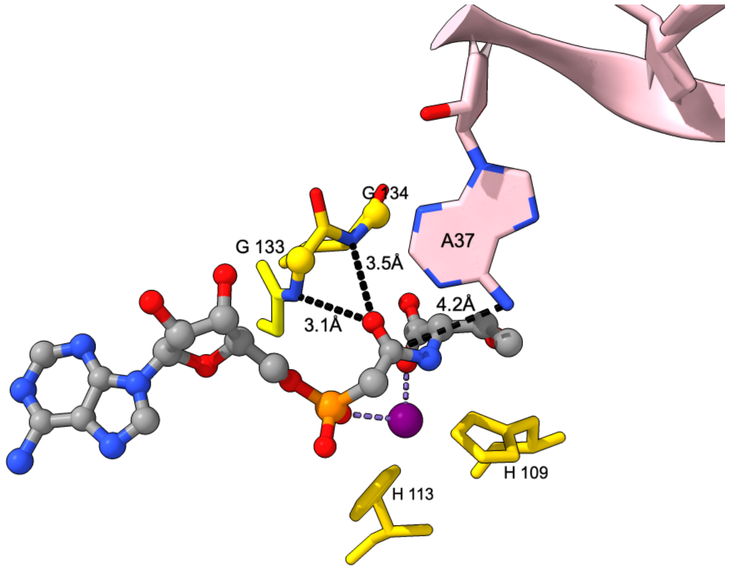


**Figure S20. Putative oxyanion hole involved in the catalytic mechanism.**
AF3 model of the OSGEP active site (same as in Fig. S10B but with a different orientation) showing the potent oxyanion hole formed by the amide nitrogens from glycine residues 133 and 134 that could stabilize the negatively charged oxygen of the tetrahedral intermediate. BK951 molecule (*i.e.* TC-AMP analog) is shown in ball-and-stick; $A_{37}$ base of tRNA in pink; $Zn^{2+}$ atom in purple and conserved OSGEP His 109 and 113 in yellow sticks.

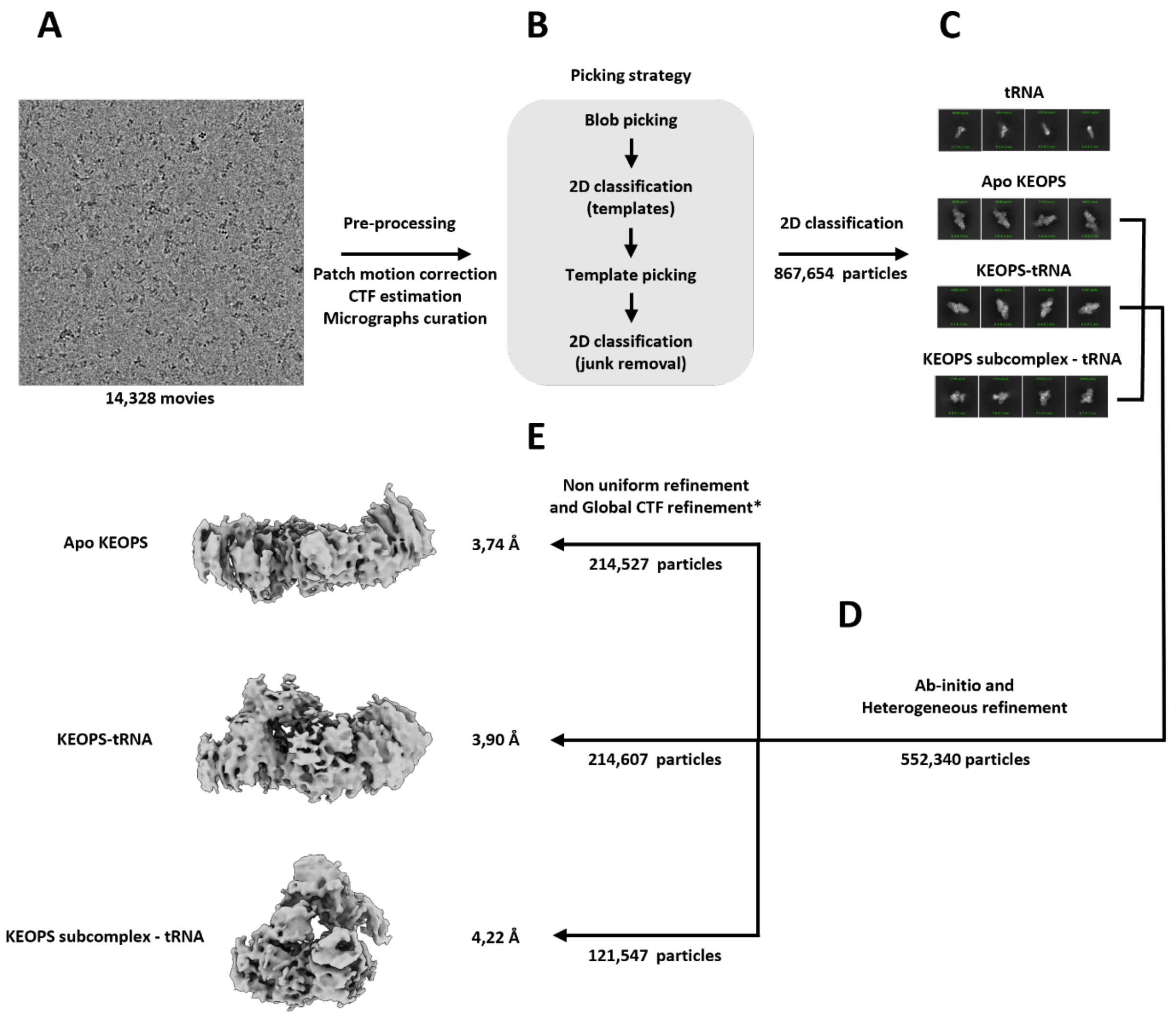


**Figure S21. Cryo-EM data processing workflow.**
All cryo-EM data processing steps, from pre-processing of the movies to final refinements of the 3D maps have been done using CryoSPARC.
(A) The movies were motion-corrected using patch motion correction and contrast transfer function estimation was done using CTFFIND4. Micrographs were screened using curate exposure according to CTF fit resolution and total full-frame motion distance.
(B) 2D templates for picking were obtained by a first automatic blob picking over the previously selected micrographs and followed by 2D classifications. To clean up the particle set obtained with template picking, two successive iterations of 2D classifications were performed and particles associated with bad classes were removed.
(C) 2D classifications of the cleaned particle set displayed particles of different sizes. Particles corresponding to tRNA alone were not used for high resolution refinements.
(D) Ab-initio reconstruction of the remaining particles yielded 3 distinct low-resolution volumes corresponding to KEOPS in complex with tRNA (KEOPS-tRNA), apo-KEOPS, and a tRNA interacting with a sub-complex of KEOPS (KEOPS subcomplex-tRNA). Heterogenous refinement was then performed to both refine the various initial models while sorting the particles between the 3 different volumes.
(E) Further refinement of the three different volumes was carried out independently using both Non-uniform refinement and Global CTF refinement. Global CTF refinement did not increase map quality or resolution of the cryo-EM map of the KEOPS subcomplex interacting with tRNA.

| ***OSGEP*** (NM-017807.4) | **cDNA** | **Protein effect** | **Gnomad v4.1.0** | **Published mutation** | **Polyphen2 Hum Var** | **SIFT** (Damaging <0.05) | **CADD** (Damaging >20) | **AlphaMissense** (Damaging >0.9) | **ACMG score*** | **Variant classification** |
|---|---|---|---|---|---|---|---|---|---|---|
| maternal allele | c.251C>T | A84V | European population: 4/1,178,836 MAF: $3.3X10^{-6.}$ 0 Homozygote | no | 0.833 possibly damaging | 0.019 | 29.7 | 0.434 ambiguous | PM1 PM2 PM3 PP2 | Likely Pathogenic |
| paternal allele | c.451G>A | E151K | European population: 20/1,179,044 MAF: $1,7X10^{-5}$ 0 Homozygote | no | 0.996 probably damaging | 0.001 | 33 | 0.986 | PM1 PM2 PP2 PP3 | Likely Pathogenic |

**Table S22. Identification of A84V/E151 pathogenic variant.**
* Variant scoring and interpretation follow the American College of Medical Genetics and Genomics guidelines[1]
**Clinical presentation:** The patient, of European descent, had exhibited global developmental delay from an early age. He later developed autistic features and microcephaly. Kidney failure with nephrotic syndrome was diagnosed at the age of 11 years
**Genetic testing results:** Targeted next-generation sequencing of a hereditary kidney disease gene panel in the patient and their parents showed that the patient carries two likely pathogenic variants in the ***OSGEP*** gene in a compound heterozygous state, which explain the patient's **GAMOS** phenotype.

[1] Sue Richards et al., 'Standards and Guidelines for the Interpretation of Sequence Variants: A Joint Consensus Recommendation of the American College of Medical Genetics and Genomics and the Association for Molecular Pathology', *Genetics in Medicine: Official Journal of the American College of Medical Genetics* 17, no. 5 (2015): 405–24, https://doi.org/10.1038/gim.2015.30.

### SUPPLEMENTARY METHODS

#### Strain, media and reagents

*Saccharomyces cerevisiae* strain IMX672 (*MAT*a *ura3-52 trp1-289 leu2-3,112 his3Δ can1Δ::cas9-natNT2*) constructed in the CEN.PK2-1C background and plasmids from the pMEL serie[2] are from Euroscarf. Liquid cultures were performed at 28°C either in rich medium (YEPD is: 1% Yeast Extract, 2% Peptone, 2% D-Glucose) or Synthetic Medium (SM is: 0.67% Yeast Nitrogen Base without amino acids, 0.2% Drop–Out Mix Complete Supplement minus uracil, tryptophan, leucine and histidine, 2% D-Glucose). When required SM was further supplemented with uracil (150 mg/L), tryptophan (75 mg/L), leucine (500 mg/L) and histidine (125 mg/L). 2% Agar were added to the recipe for plates. pW(ySua5) and pW(ΔN33Qri7p) were constructed from pFL39 backbone (ATCC). Oligonucleotides were synthetized by Eurofins. Optimization of human cDNA sequences for efficient expression in yeast were done online using JCat, the Java Codon Adaptation Tool (https://www.jcat.de/). Selection for the best CRISPR/Cas9 targeted sequence in yeast genes (20bp sequence followed by the NGG PAM sequence) and the design of the associated small guide RNA encoding DNA (120 bp centered on the selected 20 bp targeted sequence flanked with 2 stretches of 50bp necessary for homologous recombination with pMEL linear backbone) were entrusted to Yeastriction online tool (http://yeastriction.tnw.tudelft.nl). Selection for targeted sequences in human cDNA optimized sequences for CRISPR/Cas9 mediated mutagenesis was performed online using CRISPRdirect tool (http://crispr.dbcls.jp/). sgRNA-encoding DNA fragments (120 bp) and repair DNA fragments (120 bp in case of deletion and up to 500 bp for point mutations) were synthesized as dsDNA (GeneStrands). All synthetic genes were from Genscript. Preparative PCR were performed using Phusion DNA Polymerase (Thermo Scientific) for the amplification of linear pMEL backbones, sgRNA encoding DNA and repair DNA. The products of amplification were systematically purified after electrophoresis in agarose gel using GeneJET Gel Extraction Kit from Thermo Scientific prior any further use. Analytical PCR were performed using Quick-Load *Taq* 2X Master Mix according to the manufacturer's protocol (New England BioLabs) in the case of gene exchange/insertion/deletion. Small-scale preparative PCR were performed using Phusion DNA Polymerase to generate sufficient amounts of DNA for further Restriction Fragment Length Polymorphism analyses (RFLP). PCR products were purified on silica columns using GeneJET PCR Purification Kit (Thermo Scientific). Purified PCR products were submitted to the action of the suitable Fast Digest restriction enzyme (Thermo Scientific) and digestion profiles analyzed on agarose gel.

---

[2] Robert Mans et al., 'CRISPR/Cas9: A Molecular Swiss Army Knife for Simultaneous Introduction of Multiple Genetic Modifications in Saccharomyces Cerevisiae', *FEMS Yeast Research* 15, no. 2 (2015), https://doi.org/10.1093/femsyr/fov004.

5-Fluoroanthranilic Acid (FAA, Fluka) and 5-Fluoroorotic Acid (FOA, Apollo Scientific Ltd) were respectively used at 0.5g/L and 1g/L for *TRP1* and *URA3* plasmid counter-selection on Synthetic Complete Agar plates (0.67% Yeast Nitrogen Base, 0.2% Drop-out Mix Complete Supplement, 2% D-Glucose, 2% Agar). Sanger Sequencing were performed at Azenta/Genewiz.

**Expression and purification of recombinant proteins**

The vector used for *S. cerevisiae* Sua5 (ySua5) expression has been described[3]. Expression vectors for the five subunits of the hKEOPS (WT and mutants), were ordered from Genscript (Piscataway, USA). The GON7, LAGE3 and OSGEP subunits were expressed as a polycistronic construct cloned in a pET24a plasmid with an hexahistidine tag coding sequence added to the 3' end of the *hGON7* gene. The genes for the TP53RK and TPRKB subunits were cloned in pET21a and pET24b plasmids, respectively, with an hexahistidine tag coding sequence added to the 5' end of both genes.

ySua5 and hKEOPS subunits recombinant proteins were expressed in Rosetta (DE3) pLysS and BL21 (DE3) Gold *E. coli* strains (Novagen), respectively. TP53RK and TPRKB were co-expressed after cotransformation with their respective vectors.

Bacteria were grown in rich 2xYT medium until the OD600nm reached 0.5 and protein overexpression was induced by adding IPTG to a final concentration of 0.5 mM for 3 hours at 37 °C. Cells were harvested by centrifugation and resuspended in lysis buffer (20 mM Tris-Cl pH 7.5, 200 mM NaCl, 5 mM 2-mercaptoethanol). In the case of hKEOPS, cells expressing GON7, LAGE3 and OSGEP and cells expressing TP53RK and TPRKB were resuspended and mixed just before lysis to induce the formation of the complex. Cells were lysed by sonication and then centrifuged at 20,000 g for 30 minutes. Proteins were purified by Ni-NTA affinity chromatography (Qiagen) followed by size exclusion chromatography on either a Superdex™ 75 column (Cytiva) for ySua5 or a Superdex™ 200 column (Cytiva) for hKEOPS. The buffer used in the size exclusion column consisted of 20 mM Hepes pH 7.5, 200 mM NaCl, 5 mM 2-mercaptoethanol. Fractions containing pure proteins were concentrated, aliquoted, flash frozen in liquid nitrogen, and stored at -80°C.

**Construction of a humanized yeast strain for the $t^6A$ biosynthetic pathway**

Protocols were based on a cloning-free CRISPR/Cas9 approach developed in budding yeast[4]. The recipient strain IMX672, coding constitutively for *Streptomyces pyogenes* Cas9, is transformed simultaneously with three linear DNA probes (linear pMEL backbone, sgRNA encoding DNA and repair DNA). The recipe for

---

[3] Ludovic Perrochia et al., 'In Vitro Biosynthesis of a Universal t6A tRNA Modification in Archaea and Eukarya', *Nucleic Acids Research* 41, no. 3 (2013): 1953–64, https://doi.org/10.1093/nar/gks1287.

[4] Mans et al., 'CRISPR/Cas9'.

transformation follows the classical LiAc/PEG method[5], adjusting the respective quantities of the three transforming DNA probes in the transformation mix: 100ng of pMEL linear backbone, 500 ng of sgRNA encoding DNA and up to 1 µg of repair DNA for $10^8$ yeast cells. Within the cell, a double event of homologous recombination at the overlapping extremities of two of the transforming DNAs ensures the reconstitution of a stable circular plasmid from the linear pMEL plasmid backbone bearing an auxotrophic marker and the 120 bp-DNA fragment coding for the guide RNA necessary to target the endonuclease activity of the endogenous *Sp*Cas9. The generated genomic double-strand DNA break is repaired through one homologous recombination (HR) event on each side of break with each end of the repair DNA. After transformation, cells were collected and resuspended in sterile water, plated on agar medium selective for the pMEL plasmid born selection marker (*URA*3, *TRP1*, *LEU2* or *HIS3*) and grown at 28°C for 4-5 days. Selected clones were screened by PCR using a set of primers flanking the region where repair has occurred. The size of amplified products was compared to the one obtained for the unmodified strain by electrophoresis on agarose gel. Positive clones were then cured from their plasmid content either using counter-selective reagents and/or growing them on non-selective medium for several rounds. After plasmid loss confirmation, Sanger sequencing was performed to check the modification.

In the case of simultaneous insertions/deletions, we have used as many different pMEL backbones (100ng each) as different sgRNA encoding DNA (500 ng each). For multiple insertions (*hGON7*, *LAGE3*, *TP53RK* and *TPRKB* at *BUD32* and *OSGEP* at *KAE1*) repair DNA were added each at around 200 ng/kb in the transformation mix. For the simultaneous scarless deletions of *CGI121*, *GON7* and *PCC1* 1µg of each repair DNA was added.

##### *In vitro* ATPase activity assay

The ATPase activity was measured using an assay that coupled the hydrolysis of ATP to the oxidation of NADH[6] in presence of saturating amounts of unmodified tRNA substrate. All activity assays respected the initial velocity conditions[7]. The reaction was performed at 37°C in a total volume of 100 µL of 50 mM Hepes pH 8, 35 mM KCl, 3.3 mM $MgCl_2$, 10 µM $ZnCl_2$, 5 mM DTT, 4 µM human $tRNA^{Ile}_{AAU}$, 1 mM phosphoenolpyruvate, 0.7 mM NADH, 0.8 U pyruvate kinase/1.1 U lactate dehydrogenase enzymes (Sigma-Aldrich) and 3 µM hKEOPS. The reaction was initiated by the addition of 0.5 mM ATP and the

---

[5] R. Daniel Gietz and Robert H. Schiestl, 'High-Efficiency Yeast Transformation Using the LiAc/SS Carrier DNA/PEG Method', *Nature Protocols* 2, no. 1 (2007): 31–34, https://doi.org/10.1038/nprot.2007.13.

[6] M. E. Pullman et al., 'Partial Resolution of the Enzymes Catalyzing Oxidative Phosphorylation. I. Purification and Properties of Soluble Dinitrophenol-Stimulated Adenosine Triphosphatase', *The Journal of Biological Chemistry* 235 (November 1960): 3322–29.

[7] Hans Bisswanger, 'Enzyme Assays', *Perspectives in Science* 1, nos 1–6 (2014): 41–55, https://doi.org/10.1016/j.pisc.2014.02.005.

ATPase activity was monitored as the rate of absorbance decrease at 340 nm using an Infinite™ 200 PRO microplate reader (TECAN) during 1 hour.

**tRNA binding using fluorescence polarization assay**

The human $tRNA^{Ile}_{AAU}$ fluorescein labelled at its 3' end was synthesized by Horizon Discovery/Dharmacon. Binding experiments were performed in duplicates according to a published protocol[8] with 40 nM $tRNA^{Ile}_{AAU}$-3'Fl and 0 to 1.6 µM hKEOPS in 50 mM Hepes pH 8, 35 mM KCl, 5 mM DTT in a final volume of 200 µL. After 30 min incubation at room temperature, the mixtures were transferred in a 96-well flat bottom black plate (Greiner) and polarization fluorescence were recorded using an Infinite™ 200 PRO microplate reader (TECAN) (excitation and emission wavelengths, 485 nm and 535 nm respectively). $K_D$ values were estimated by fitting the binding curves using the one-site rectangular hyperbola equation of the Graphpad Prism 10.6 software.

**T7 *in vitro* transcription and purification of tRNA.**

The double-stranded DNA template used for the transcription of human $tRNA^{Ile}_{AAU}$ by T7 RNA polymerase was prepared by PCR. Two oligonucleotides complementary at their 3'ends were used as primers and templates for PCR reaction by the Phusion™ High-Fidelity DNA Polymerase (Thermo Scientific). The oligonucleotide sequences were [U2'OMe][G2'OMe]GTGGCCCGTACGGGGATCGAACCCGCGACCT TGGCGTTATTAGCACCACGCTCTAAC (forward primer) with two 2'O methylated nucleotides at the 5' end to optimize the T7 transcription stop[9] and TAATACGACTCACTATAGGCCGGTTAGCTCAGTTGG TTAGAGCGTGGTGCTAATAAC (reverse primer) with T7 promoter sequence at the 5' end to perform *in vitro* T7 RNA transcription. After phenol-chloroform extraction and ethanol precipitation, the PCR product (60 µg) was used as a template for T7 run-off transcription reaction in a final volume of 5 mL of 40 mM Tris-HCl pH 8.0, 1 mM spermidine, 5 mM DTT , 0.01% (w/v) Triton X100, 20 mM $MgCl_2$, 4 mM ATP/GTP/CTP/UTP, 200 µg recombinant T7 RNA polymerase (homemade) at 37 °C for 4 h. Transcription was stopped by adding 50 mM EDTA, and transcript was purified by preparative electrophoresis on TBE-urea polyacrylamide gel (10% Acrylamide:Bis-acrylamide 19:1). The tRNA was visualized in the gel by UV shadowing and the corresponding band was cut out. The band was transferred into a dialysis tube (Spectra/por MWCO 6-8 kDa) containing 4 mL TBE 0.25x and electroeluted by using an horizontal

[8] Marie-Claire Daugeron et al., 'A Paralog of Pcc1 Is the Fifth Core Subunit of the KEOPS tRNA-Modifying Complex in Archaea', *Nature Communications* 14, no. 1 (2023): 526, https://doi.org/10.1038/s41467-023-36210-y.

[9] C. Kao et al., 'A Simple and Efficient Method to Reduce Nontemplated Nucleotide Addition at the 3′ Terminus of RNAs Transcribed by T7 RNA Polymerase', *RNA* 5, no. 9 (1999): 1268–72, https://doi.org/10.1017/S1355838299991033.

electrophoresis tank as described [10]. The electro-eluate was subjected to ethanol precipitation and the tRNA pellet was resuspended in DEPC-treated water (5 mL) and desalted by 3 concentration/dilution cycles by centrifugating at 2500 g in Vivaspin 20 MWCO 10 kDa (Sartorius). The tRNA was refolded by successive incubations at 80°C for 3 min and on ice for 30 min.

**tRNA extraction, purification and hydrolysis**

Total yeast tRNAs were extracted and purified from actively growing cells at an $OD_{600nm}$ of about 1 (30mL of culture at $3.10^7$cells/mL) using phenol-induced cell permeabilization, LiCl selective precipitation and subsequent ion exchange chromatography purification on an AXR-80 column (Nucleobond, Macherey-Nagel) following the manufacturer's instructions. 10µg samples of purified tRNAs were enzymatically hydrolyzed into ribonucleosides using P1 Nuclease, Phosphodiesterase and Alkaline Phosphatase according to previously published protocols [11]. Samples were loaded on a 10 kDa Microcon centrifugal unit to remove enzymes and the recovered filtrate that contains nucleosides (flow-through) was finally dried under vacuum.

**LC-HRMS quantification of A and** $t^6A$ **after tRNA digestion**

The nucleosides were quantitated by liquid chromatography coupled to a high-resolution mass spectrometer (LC-HRMS). Nucleosides were identified according to their specific elution times and exact *m/z* values. The experiments were performed with a LC 1260 Prime coupled to a QTOF 6546 (Agilent Technologies, Waldbronn, Germany). The column (HSS T3, 150 mmm – 2.1 mm × 2.7 µm, Waters, USA) was chosen for an efficient separation of the canonic and modified nucleosides without the use of any buffer, such as ammonium formiate, in the mobile phase. The solvent A was water + 0.1% formic acid (Sigma-Aldrich, Saint-Quentin Fallavier, France) and solvent B acetonitrile (LC grade, J.T. Baker). The gradient started at 100% of A to 50% in 8 min then 100% of solvent B during 2 min and finally back to the initial conditions. The injected volume was fixed at 2 µL. For electrospray (ESI) analysis, mass spectra were recorded in positive ion mode with the following parameters: gas temperature 325°C, drying gas flow rate 10 l $min^{-1}$, nebulizer pressure 30 psi, sheath gas temperature 400°C, sheath gas flow rate 10 l $min^{-1}$, capillary voltage 3500 V, nozzle voltage 500 V, fragmentor voltage 110 V, skimmer voltage 45 V, Octopole 1 RF Voltage

[10] Sara Lopez-Gomollon and Francisco Esteban Nicolas, 'Purification of DNA Oligos by Denaturing Polyacrylamide Gel Electrophoresis (PAGE)', in *Methods in Enzymology*, vol. 529 (Elsevier, 2013), https://doi.org/10.1016/B978-0-12-418687-3.00006-9.

[11] Christelle Arrondel et al., 'Defects in t6A tRNA Modification Due to GON7 and YRDC Mutations Lead to Galloway-Mowat Syndrome', *Nature Communications* 10, no. 1 (2019): 3967, https://doi.org/10.1038/s41467-019-11951-x; Kathrin Thüring et al., 'Analysis of RNA Modifications by Liquid Chromatography–Tandem Mass Spectrometry', *Methods* 107 (September 2016): 48–56, https://doi.org/10.1016/j.ymeth.2016.03.019.

750 V. For ESI, internal calibration was achieved with two calibrants purine and hexakis (1 h,1 h,3 h-tetrafluoropropoxy) phosphazene (*m/z* 121.0509 and *m/z* 922.0098) providing a high mass accuracy better than 3 ppm. First of all, standards, including A (Sigma-Aldrich) and $t^6A$ (BioLog Life Science Institute, Bremen, Germany) were injected to determine the elution times and calibration curves from 10 to 500 nM injected in triplicate. For quantification, the nucleoside samples were diluted into different concentrations (as determined by the OD at 254 nm) to enable quantification on either A or $t^6A$ based on their relative concentrations.

**Michaelis-Menten saturation curves of $I14F^{OSGEP}$, $V107M^{OSGEP}$, $I111T^{OSGEP}$ and $R325Q^{OSGEP}$.**

3 µM of each GAMOS mutant activity have been assayed in parallel with the same concentration (3 µM) of WT KEOPS by using 5 increasing concentrations of $htRNA^{Ile}{}_{AAU}$ substrate ranging from 2.5 µM to 20 µM. Saturation curves have been fited with Graphpad Prism 10.6 software using the quadratic following equation in order to account for absence of substrate excess :

$$Vi = \frac{k_{cat}\left( [E_t] + [S_0] + K_M - \sqrt{([E_t] + [S_0] + K_M)^2 - 4[E_t][S_0]} \right)}{2}$$

Catalitic constant $k_{cat}$ and Michaelis constant $K_M$ have been obtained by non-linear regression and catalytic efficiency (or specificity constant) $k_{cat}/K_M$ have been calculated. The ratio $(k_{cat}/K_M)^{mutant}/(k_{cat}/K_M)^{WT}$ is given in parenthesis in Fig S15.
